# Biomolecular Condensation and L-Cysteine Signaling Activates Dormant Protease Activity of Papain Droplets

**DOI:** 10.64898/2026.06.29.735447

**Authors:** Saurabh Gupta, Brijeshwar Singh, Prashant Kodgire, Tushar Kanti Mukherjee

## Abstract

Proteases are an important class of proteolytic enzymes having great importance in both basic science and industrial applications. While cells tightly regulate the spatio-temporal activity of different proteases for cellular homeostasis, mis-regulation often leads to adverse effects. In this context, the protease activity of papain and its activation by L-cysteine is poorly understood in the literature. Herein, we discover that the protease activity of papain can be effectively regulated via a spontaneous liquid-liquid phase separation (LLPS) pathway. We show that papain undergoes biomolecular condensation via spontaneous LLPS under macromolecular crowding through the involvement of intermolecular hydrophobic interactions. Secondary structure analyses revealed a compact conformation of phase-separated papain with increased *α*-helix content. Although native free papain is found to be active towards synthetic and protein substrates, the proteolytic digestion produces heterogeneous peptide aggregates. In contrast, we found that papain droplets remain dormant toward protein digestion due to the disulfide linkage of the active cysteine residue (Cys-25) in its compact conformational state. More importantly, we show that the protease activity of phase-separated papain can be reactivated in the presence of L-cysteine to produce uniform soluble peptide fragments. Our findings indicate that although disulfide linkages are not necessary for the phase separation of papain, upon phase separation, intermolecular interactions between phase-separated papain result in the formation of disulfide linkages involving active Cys-25 residues. The present discovery has tremendous technological importance to boost the efficacy of meat tenderization in the food industry.

## INTRODUCTION

Hydrolysis of a peptide bond is an energetically favorable reaction, but an extremely sluggish process.^1^ Cells utilize proteases or proteolytic enzymes to hydrolyse the peptide bonds present within proteins with high efficiency and selectivity.^2,3^ These proteases are involved in many important physiological processes, including protein turnover, digestion, blood coagulation, wound healing, cell signaling, cell growth, immune response, and apoptosis.^2^ Uncontrolled or unregulated proteolysis can lead to many adverse disease states, including emphysema, stroke, cancer, Alzheimer’s, inflammation, and arthritis.^4^ Therefore, cells must tightly regulate the spatio-temporal activity of these proteases for cellular homeostasis. Understanding the inherent catalytic cycles and their regulatory mechanisms is of great importance in cellular biology as well as in industrial applications.^5,6^

Proteases can be classified into six major groups based on their catalytic mechanism: cysteine, aspartic, serine, threonine, glutamic, and metallo-proteases.^7^ Among these, papain is the most studied cysteine protease and is a member of the C1 family.^8–10^ Papain is a monomeric polypeptide with a molecular weight of 23.4 kDa and its active form consists of 212 amino acid residues with three disulfide bridges between Cys22–Cys63, Cys56–Cys95, and Cys153–Cys200.^11^ The active site of papain consists of a catalytic triad having residues Cys-25, His-159, and Asn-175. Although papain primarily breaks peptide bonds present in basic amino acids, especially arginine, lysine, and phenylalanine, it cleaves a wide variety of peptide bonds, indicating a broad specificity.^12,13^ The Cys-25 residue acts as a primary nucleophile, and the His-159 residue facilitates the nucleophilic attack at the carbonyl groups of amide bonds by abstracting the proton from the Cys-25.^2,14,15^ This nucleophilic attack at the carbonyl groups of amides results in the formation of a tetrahedral intermediate which is stabilized by an oxyanion hole through several hydrogen bond donors near the active site. On the other hand, the Asn-175 residue at the active site stabilizes the tetrahedral transition state. This enables papain to efficiently catalyze the cleavage of peptide bonds. Although the mechanistic aspect of the protease activity of papain is well established in the literature, the crucial role of L-cysteine (L-Cys) on the activation of papain activity is poorly understood and controversial. Earlier, it was proposed that the structure of the major fraction of inactive papain contains a mixed disulfide between active Cys-25 of papain and cysteine.^16–19^ It was shown that activation of inactive papain is promoted by additives such as thiols, cyanide, or sodium borohydride. In contrast, Pihl and co-workers observed that the papain used was active without any special pre-treatment or additives.^20^ Their kinetic data contradict the earlier hypothesis that the active Cys-25 residue is bound in the form of an internal thiester/disulfide bond. Through their detailed kinetic measurements it was shown that the activation by cystein was associated with an increase in *K*_m_ value via enhancing the rate of cleavage of the enzyme-substrate complex. Moreover, most of these previous studies were performed under dilute buffer conditions, which lack physiologically relevant heterogeneous crowded environments. To fill this gap and better understand the role of L-cysteine activation, we have studied the protease activity of papain under cell mimicking crowded environments under physiological conditions in the absence and presence of L-cysteine.

The heterogeneous and crowded cellular milieu significantly alters the physicochemical properties of biomolecules^21–23^ and modulates various fundamental biochemical processes, including enzymatic activity,^24–28^ protein folding,^29–31^ and DNA replication.^32^ Our recent findings on enzymatic activities in crowded environments revealed a highly efficient hidden pathway via liquid-liquid phase separation (LLPS), which is completely ignored in the literature.^33,34^ It is now a well established phenomenon that macromolecular crowding promotes liquid-liquid phase separation (LLPS) of many intrinsically disordered^35–39^ as well as ordered proteins^40–43^ via enhancing multivalent intermolecular interactions. These interactions are mainly weak non-covalent attractive forces that stabilize the condensates/droplets in aqueous dispersion. Notably, macromolecular crowding elevates these intermolecular protein-protein interactions via the excluded volume effect by lowering the critical protein concentration needed for LLPS. Previously, we observed that phase-separated droplets of functional enzymes spatio-temporally regulate the enzymatic activity in the absence and presence of different signaling mechanisms.^33,34^ In this context, no systematic study on the protease activity of papain and the role of L-cysteine activation has been undertaken under crowded milieu. To fill this gap and motivated by our previous findings, we have undertaken the present study to understand the protease activity of papain and the regulatory role of L-cysteine under heterogeneous and crowded environments. Herein we have demonstrated a unique and hidden regulatory role of L-Cys behind the highly efficient protease activity of papain via the LLPS pathway using both synthetic and natural protein substrates (Figure 1a). Our findings reveal that papain undergoes spontaneous phase separation under inert macromolecular crowding to yield uniform spherical droplets. We show that precise L-cysteine signaling is essential to transform these dormant droplets into active droplets, which efficiently sequester protein substrates for efficient digestion. To the best of our knowledge, this is the first report to demonstrate the hidden regulatory role of L-Cys behind the proteinase activity of papain via the LLPS pathway (Figure 1a).

**Figure 1.**
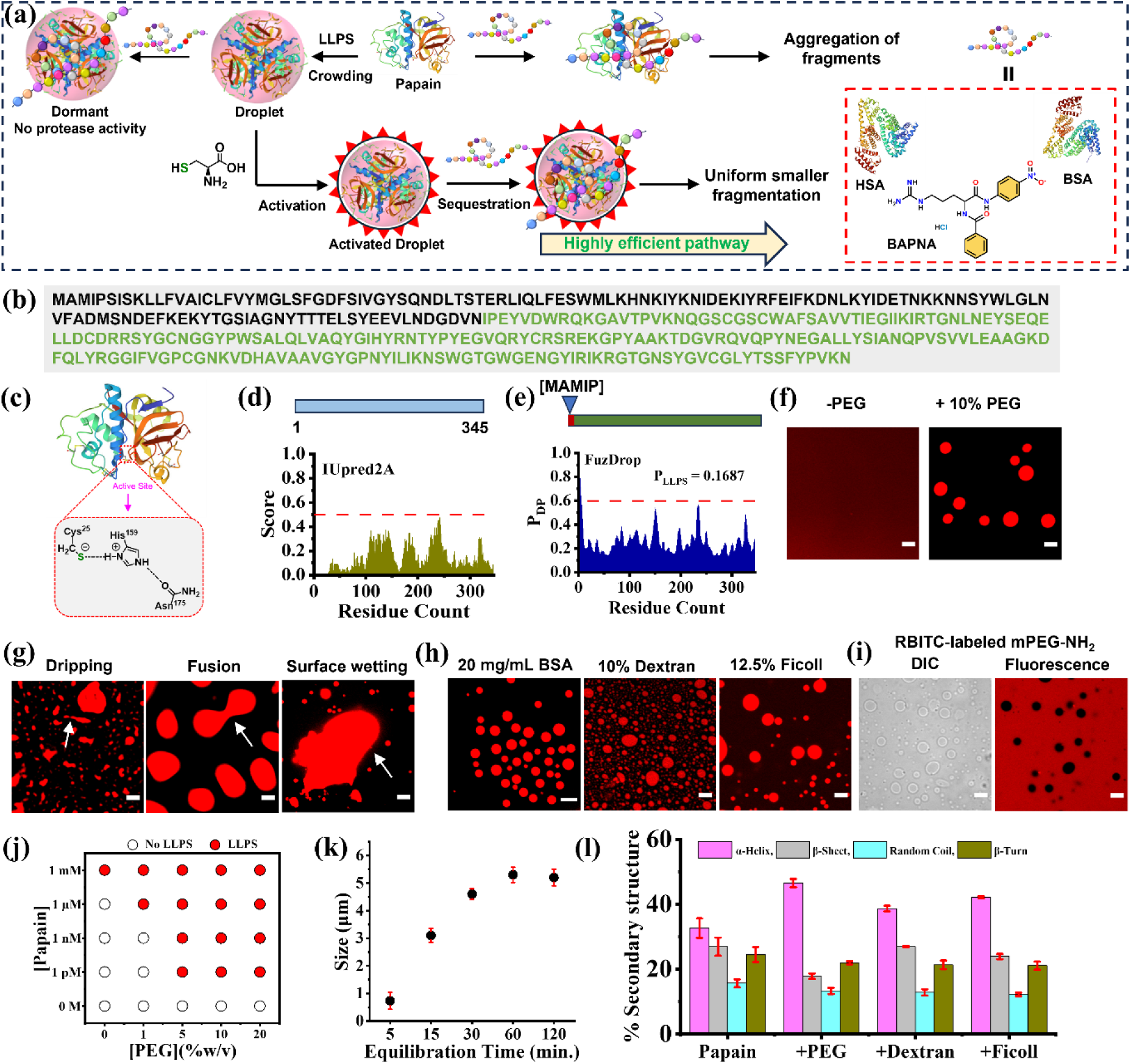
(a) Schematic showing spatio-temporal regulation of protease activity of papain via LLPS and L-Cys activation. (b) Primary amino acid sequence of the proenzyme form of papain obtained from the Protein Data Bank (PDB entry 9PAP). The primary sequence of active papain is highlighted in green color. (c) Crystal structure and active site of papain obtained from *Carica Papaya* (PDB entry 9PAP). Bioinformatics analyses using (d) SMART (upper panel) algorithm for LCDs and IUPred2A (lower panel) for disorder regions, and (e) FuzDrop algorithm to predict the probability of LLPS. (f) Confocal images of 1.66 μM RBITC-labeled papain in the absence and presence of 10% PEG-8000. (g) Confocal images of papain droplets showing dripping, fusion, and surface wetting phenomena. (h) Droplet formation in the presence of 20 mg/mL BSA, 10% dextran 70, 12.5% Ficoll 400. (i) Confocal images of unlabeled papain in the presence of RBITC-labeled 10% mPEG-NH_2_ (MW 5000). (j) Phase diagram of papain as a function of PEG concentrations. (k) Variation of mean droplet size as a function of equilibration time in the presence of 10% PEG 8000. (l) Variation in the secondary structure contents of papain in the absence and presence of different crowders estimated from deconvoluted FTIR spectra. All samples were prepared in 50 mM phosphate-citrate buffer (pH 6.0) at 37 °C. Scale bars correspond to 5 µm.

## RESULTS AND DISCUSSION

### Macromolecular Crowding Promotes LLPS of Papain

The inactive proenzyme form of papain consists of 345 amino acid residues, and among these, only 212 amino acids are present in the active and mature papain (Figure 1b). The crystal structure of papain revealed the substrate binding active site having amino acid residues of Cys-25, His-159, and Asn-175 (Figure 1c). The primary sequence analyses of papain using SMART^44^ and IUpred2A^45^ algorithms revealed neither any LCDs nor any IDRs in the amino acid sequence (Figure 1d). In addition, sequence-based prediction using the FuzDrop algorithm revealed a very low value of the propensity of papain to form spontaneous droplets (*p*_DP_ < 0.6) and an overall LLPS propensity (*p*_LLPS_) of 0.1687 (Figure 1e).^46^ With these initial bioinformatics findings, we performed confocal laser scanning microscopy (CLSM) with 1.66 µM of RBITC-labeled papain in the absence and presence of different synthetic (PEG, dextran, and Ficoll) and protein (BSA) crowders. In the absence of crowders, the CLSM image of RBITC-labeled papain revealed diffused background fluorescence signals without any specific structures (Figure 1f). In contrast, distinct spherical structures with localized red emission were observed in the presence of 10% (w/v) PEG 8000. These spherical assemblies exhibited characteristic dripping, fusion, and surface wetting phenomena (Figure 1g), suggesting that these assemblies are liquid-like droplets or condensates as reported previously.^33–43^ Similar liquid-like droplets with distinct red emission were also observed in the presence of 20 mg/mL BSA as protein crowder along with 10% (w/v) dextran 70, and 12.5% (w/v) Ficoll 400 as inert synthetic crowders (Figure 1h), suggesting that the droplet formation is a general phenomenon for papain in the presence of inert crowders. Notably, the mean size of these droplets increased appreciably with an increase in papain concentration at a given concentration of 10% PEG 8000 (Figure S1), suggesting enhanced protein-protein interactions and a higher rate of fusion events at higher protein concentrations. Moreover, no liquid-to-solid phase transition was observed for these papain droplets over a period of 30 days (d 30), suggesting that the phase-separated papain remains conformationally stable against irreversible protein aggregation (Figure S2).

In general, proteins form liquid-like droplets via LLPS either via homotypic or heterotypic interactions. To know whether the phase separation of papain is due to homotypic or heterotypic interactions, we performed the phase separation assay with unlabeled papain and RBITC-labeled 10% mPEG-NH_2_ (MW 5000) as a crowder (Figure 1i). While the phase contrast image revealed the presence of well-dispersed spherical droplets, the CLSM image showed that the labeled PEG was completely excluded from the droplet phase as no detectable fluorescence signals were observed from inside the droplets (Figures 1i and S3). This authenticates that the present LLPS of papain is a homotypic phase separation and occurs only due to homotypic protein-protein interactions between papain molecules. Notably, droplet formation was observed in the absence of crowders only at a very high concentration (1 mM) of papain (Figures 1j and S4). Moreover, the critical concentration of papain needed for phase separation decreased with an increase in the PEG concentrations (Figures 1j and S4), suggesting enhanced protein-protein interactions even at lower protein concentration due to the excluded volume effect at high crowder concentration. The mean size of these droplets increased noticeably with equilibration time and saturated beyond 1 h of equilibration (Figures 1k and S5). This observation can be explained by considering spontaneous fusion and Ostwald ripening of liquid-like droplets in aqueous suspension.^42^ Taken together, these findings authenticate that macromolecular crowding is essential to promote LLPS of papain via homotypic protein-protein interactions in aqueous buffer to yield biomolecular droplets. To know whether native papain undergoes any conformational alteration during crowding-induced phase separation, we thoroughly analyzed the secondary structure of papain before and after the phase separation in the presence of different crowders using FTIR measurements. The FTIR spectral analysis of native papain reveals 33% *α*-helix, 27% *β*-sheet, 16% random coil, and 24% *β*-turn contents, which matches well with the previous report.^47^ However, in the presence of 10% PEG, the *α*-helix content of papain increased to 47% with concomitant decrease in the *β*-sheet, random coil, and *β*-turn contents to 18, 13, and 22%, respectively (Figures 1l and S6). A similar trend was also observed in the presence of 10% dextran and 12.5% Ficoll as crowders (Figures 1l, S7, and S8). These findings suggest that papain attains a more compact conformational state upon phase separation in the presence of different crowders through the involvement of homotypic protein-protein interactions. Next, we sought to address the nature of these soft protein-protein interactions via several control experiments.

### Intermolecular Interactions Behind LLPS

Soft protein-protein interactions are known to be influenced by pH and temperature of the medium.^33–43^ To know the influence of pH and temperature on the LLPS of papain, we performed LLPS assays of papain by varying the solution pH and temperature from 2.0–11.0 and 4–90 ℃, respectively (Figure 2a,c). The CLSM images revealed stable droplet architecture in the pH range of 3.0–11.0; however, they were unstable at a lower acidic pH of 2.0 (Figures 2a and S9).

**Figure 2.**
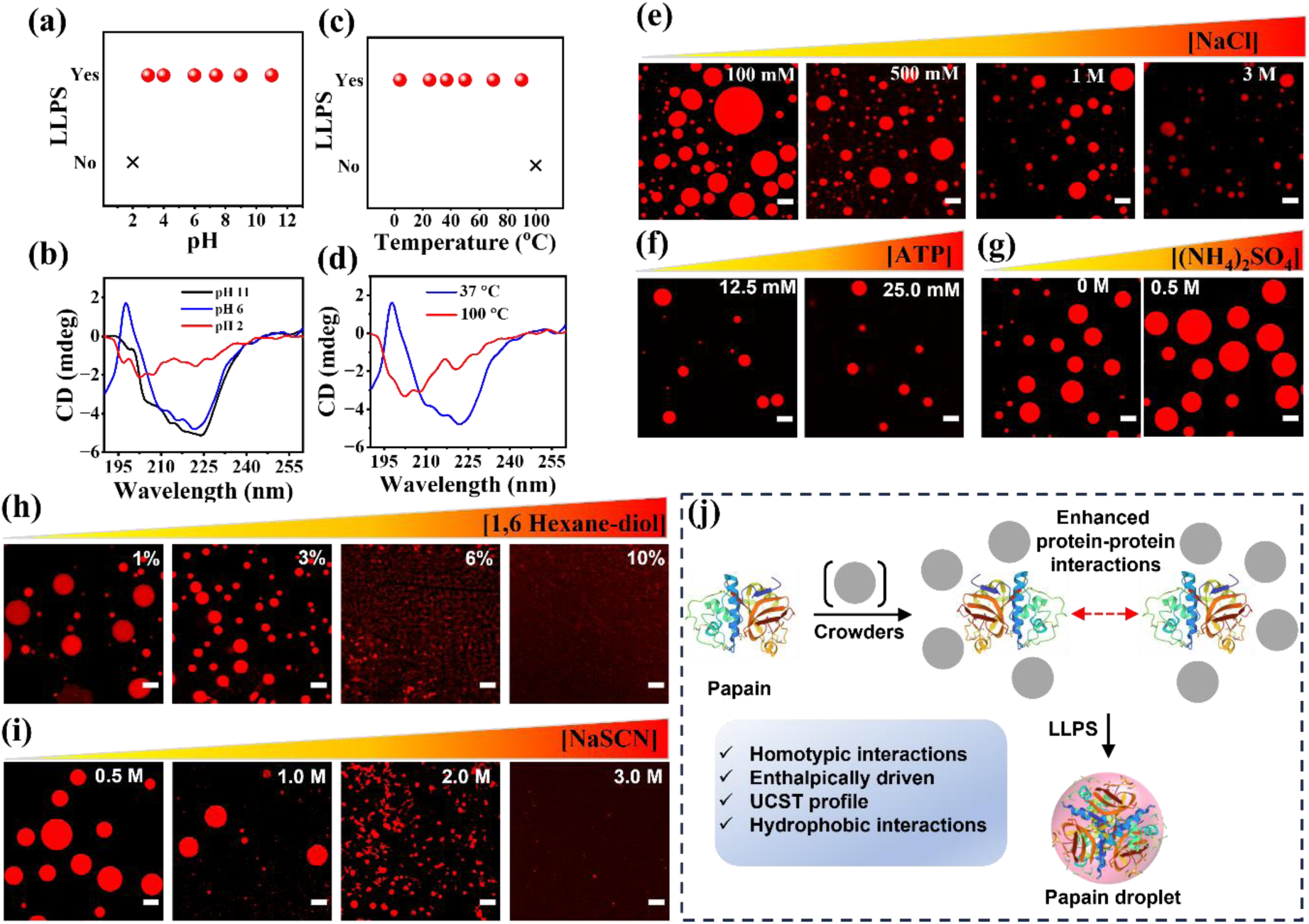
Effect of pH and temperature on the (a, c) phase separation and (b, d) secondary structures of papain in the presence of 10% PEG 8000. Confocal images of RBITC-labeled papain droplets as a function of different concentrations of (e) NaCl, (f) ATP, (g) (NH_4_)_2_SO_4_ (h) 1,6-hexanediol, (i) NaSCN. (j) Schematic representation of the crowding-induced LLPS of papain via homotypic protein-protein interactions. Samples were prepared in 50 mM phosphate-citrate buffer at pH 6.0 and incubated at 37 °C for 1 h. Scale bars correspond to 5 µm.

Notably, the droplet size decreased appreciably at pH 3.0 (Figure S9), signifying weakening of protein-protein interactions at lower acidic pH. Although the reported isoelectric point (pI) of papain is ∼8.75,^14,48^ we observed spontaneous LLPS up to pH 3.0. This signifies that the overall charge of papain has a negligible impact on the feasibility of LLPS, rather short-range interactions between peptide residues play a critical role in the LLPS of papain. The far-UV CD data revealed significant conformational destabilization at pH 2.0 compared to pH 6.0 and 11.0 (Figure 2b). Therefore, our findings reveal that papain undergoes spontaneous LLPS in a broad range of pH from 3.0–11.0; however, conformationally altered papain at a lower acidic pH of 2.0 failed to undergo LLPS due to the lack of favorable protein-protein interactions.

Next, we investigated the effect of temperature on the feasibility of LLPS of papain. Phase separation assays were performed with 1.66 µM papain in the presence of 10% PEG 8000 by varying the temperature in the range of 4–100 °C under CLSM (Figures 2c and S10). Papain droplets were found to be stable in the temperature range between 4–90 °C; however, phase separation and subsequent droplet formation were suppressed at 100 °C (Figures 2c and S10). The mean droplet size decreases appreciably to 1.18 ± 0.27 µm at 90 °C from that of 3.74 ± 1.30 µm at 37 °C. The far-UV CD measurements revealed significant alteration of the native conformation of papain at an elevated temperature of 100 °C compared to that at 37 °C (Figure 2d). Notably, the unstructured CD spectrum of papain at 100 °C resembled with that observed at a lower acidic pH of 2.0 in the presence of 10% PEG 8000. This suggests that conformationally denatured papain at acidic pH (pH ≤ 2.0) and elevated temperature (T ≥ 100 °C) fails to undergo LLPS, possibly due to the lack of favorable intermolecular protein-protein interactions. Previously, it has been shown that proteins can either display upper critical solution temperature (UCST)^35^ or lower critical solution temperature (LCST)^37^ or both^49^ depending on the intermolecular interaction forces. While the UCST-based phase separations of biomolecules are primarily driven by enthalpically favorable (Δ*H* < 0) intermolecular protein-protein interactions,^33,40,41^ the LCST-based phase separations are mainly driven by entropy.^37^ The present UCST profile of papain condensation indicates that the LLPS of papain is primarily driven by enthalpically favorable protein-protein interactions under macromolecular crowding. It should be noted that the essential role of macromolecular crowding is to promote favorable intermolecular protein-protein interactions, which facilitate overcoming the unfavorable entropy loss associated with the demixing process (Figure 2j). Notably, biomolecular condensation is thermodynamically favorable (ΔG < 0) only when the enthalpy gain via multivalent intermolecular interactions overcomes the entropy of mixing according to the Flory–Huggins theory.^50,51^ Taken together, our present data indicate that papain undergoes LLPS through enthalpically driven protein-protein interactions in a broad range of pH (3.0–11.0) and temperature (4–90 °C) under macromolecular crowding.

Next, to decipher the nature of the intermolecular protein-protein interactions behind the phase separation of papain, we performed a series of control phase separation experiments in the presence of different additives that perturb various protein-protein interactions. In general, biomolecules undergo LLPS through the involvement of various non-covalent intermolecular interactions, including electrostatic, hydrophobic, π–π, cation–π, and/or hydrogen bonding. Initially, to check whether electrostatic interactions have any role in the phase separation of papain, we varied the ionic strength of the phosphate-citrate buffer (pH 6.0) by changing the NaCl concentration from 100 mM to 3 M. CLSM images revealed that, irrespective of the NaCl concentrations, phase separation was observed even at a very high NaCl concentration of 3 M (Figure 2e). This indicates that the phase separation of papain is independent of the ionic strength of the medium, and electrostatic interactions play a negligible role in the LLPS of papain. This argument gains support from a control experiment performed in the presence of different concentrations of ATP, which acts like an electrostatic disruptor. CLSM images revealed intact, well-dispersed spherical droplets in the presence of 12.5 and 25.0 mM ATP (Figure 2f), suggesting a lack of any electrostatic interactions between papain molecules during the LLPS process. On the other hand, the addition of a kosmotropic salt, ammonium sulfate (NH_4_)_2_SO_4_, into the buffer favored the condensation of papain, as revealed by the formation of larger-sized droplets in the CLSM images (Figures 2g and S11). This suggests a possible role of hydrophobic interactions between papain molecules in the condensation process. To further authenticate the role of hydrophobic interactions for the condensation of papain, we performed LLPS assays in the presence of varying concentrations of 1,6-hexanediol and sodium thiocyanate (NaSCN), which are known to disrupt the hydrophobic protein–protein interactions. CLSM images revealed well-dispersed droplets of papain in the presence of 1 and 3% of 1,6-hexanediol; however, droplet formation was inhibited at higher concentrations (6 and 10%) of 1,6-hexanediol (Figure 2h), suggesting that the LLPS of papain is primarily driven by hydrophobic interactions. Similarly, condensation of papain was found to be completely hindered in the presence of 3.0 M of NaSCN (Figure 2i). Altogether, these findings authenticate the crucial role of intermolecular hydrophobic interactions between the papain molecules in the LLPS of papain under macromolecular crowding (Figure 2j).

### Protease Activity of Free and Phase-Separated Papain toward Synthetic Substrate

Upon establishing the crowding-induced spontaneous LLPS of papain through multivalent hydrophobic interactions, we next examined and compared the protease activity of free papain with that of phase-separated papain inside the droplet. Previously, our group illustrated that LLPS of functionally active enzymes provides an alternative and highly efficient pathway to modulate their biocatalytic efficacy.^33,34^ We envisaged that the protease activity of phase-separated papain with altered conformation might differ significantly from that of free papain and may open up a new avenue for its future applications. Papain, being a cysteine protease exhibits a broad substrate scope and can digest a wide variety of substrates, which leads to an elevated level of cook loss.^52,53^ To explore the protease activity, we utilized a synthetic substrate, *N_α_*-benzoyl-arginine-*p*-nitroanilide (BAPNA), which has been extensively utilized as a model substrate to evaluate the activity of papain and related protease enzymes (Figure _3a__).12,54–58_

**Figure 3.**
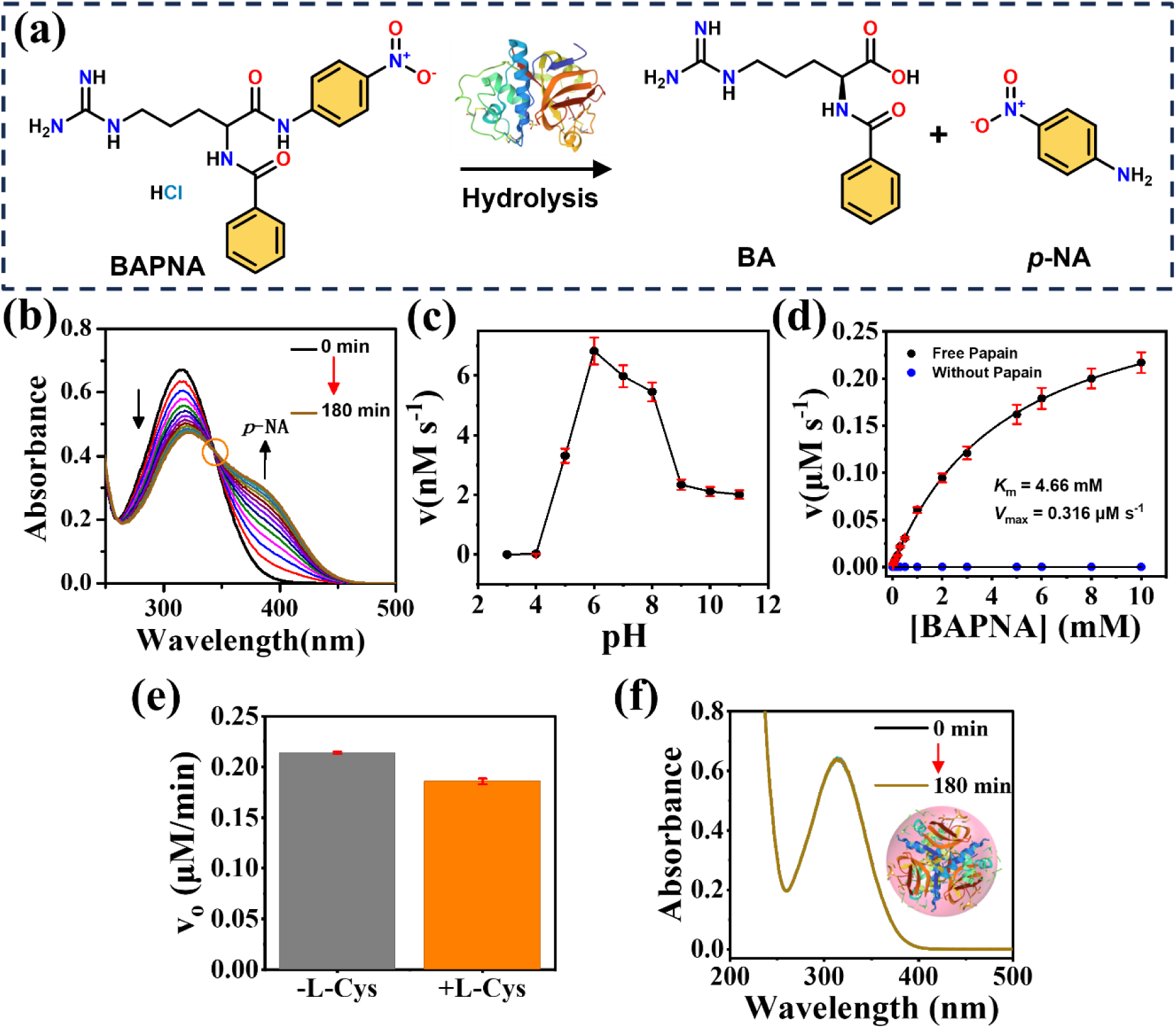
(a) Schematic showing the protease activity of papain using BAPNA substrate. (b) Time dependent UV-vis spectra of 50 µM BAPNA in the presence of 1.66 µM papain at 15 min intervals for 180 min. (c) Effect of pH on the initial rate of hydrolysis of 100 µM BAPNA by 1.66 µM papain. (d) Michaelis-Menten plot of BAPNA hydrolysis in the absence and presence of papain at pH 6.0. (e) Plot of initial velocity for the hydrolysis of 50 µM BAPNA by free papain in the absence and presence of 5 mM L-Cys. (f) Time dependent UV-vis spectra of 50 µM BAPNA in the presence of papain droplets. Samples were prepared in a 50 mM phosphate-citrate buffer (pH 6.0) at 37 °C.

The hydrolysis of BAPNA in the presence of papain can be conveniently monitored using UV-vis spectroscopy by recording the characteristic spectral feature of *p*-nitroaniline (*p*-NA) at a wavelength (*λ*_max_) of 400 nm (Figure 3a,b). Samples were prepared with 1.66 µM papain in phosphate-citrate buffer at pH 6.0 and equilibrated at 37 °C for further investigations. BAPNA alone in phosphate-citrate buffer at pH 6.0 showed an absorption band at 316 nm (Figure 3b). However, the addition of 1.66 µM papain resulted in the lowering of the original absorbance at 316 nm with concomitant appearance of a new absorption band centred at 400 nm upon 180 min of reaction (Figure 3b). The UV-vis kinetic spectra also reveal an isosbestic point at 343 nm, suggesting stoichiometric conversion of BAPNA to *p*-NA in the presence of papain. The enzymatic rates of papain at different experimental conditions were monitored by recording the changes in the absorbance at 400 nm as a function of reaction time. Notably, pH dependent hydrolysis of BAPNA by papain revealed maximum protease activity at pH 6.0 (Figure 3c). The rate of hydrolysis of BAPNA in the presence of papain follows a typical Michaelis–Menten behavior as a function of BAPNA concentrations in the range of 0–10 mM (Figure 3d). No such conversion of BAPNA to *p*-NA was observed in the absence of papain (Figures 3d and S12), highlighting the catalytic role of papain in hydrolyzing the peptide bond. The kinetic data were fitted with the classical Michaelis–Menten equation to determine the maximum velocity (*V*_max_) and Michaelis constant (*K*_m_). The estimated values of *V*_max_ and *K*_m_ for the hydrolysis of BAPNA by papain were found to be 0.316 µM s^-1^ and 4.66 mM, respectively. The associated turnover number (*k*_cat_) was calculated to be 0.19 s^-1^ at 37 °C and pH 6.0. These kinetic parameters closely match the previous reports of BAPNA hydrolysis by papain.^12,55,56^ Taken together, these findings authenticate that free papain is active toward the synthetic substrate BAPNA at pH 6.0 and 37 °C. Notably, the addition of L-Cys showed no significant impact on the hydrolysis of BAPNA in the presence of free papain under similar experimental conditions (Figure 3e), except for a marginal decrease. This indicates that free papain is active in its native state, and the active Cys-25 residue remains in its free thiol form.

Next, we checked the activity of the papain droplet toward the hydrolysis of BAPNA under similar experimental conditions. For this, we removed the free PEG present in the dilute phase by centrifugation, and the isolated condensed phase was redispersed in phosphate-citrate buffer at pH 6.0. Notably, no noticeable change in the UV-vis absorption spectra of BAPNA was observed over a period of 180 min in the presence of papain droplets (Figure 3f). The absence of the characteristic absorption band at 400 nm of *p*-NA suggests that phase-separated papain inside the droplet phase is incompetent to hydrolyse the peptide bond of BAPNA. This intriguing observation can be explained by considering two possible scenarios. Firstly, the lack of partition of BAPNA inside the droplet phase may result in no proteinase activity. To check this, we estimated the partition coefficient of BAPNA into the papain droplet using UV-vis spectroscopy. The estimated partition coefficient of BAPNA into papain droplets is found to be 3.60 ± 0.62, suggesting that BAPNA has a high affinity toward papain droplets. Secondly, conformational alteration-induced inactivation of papain inside the droplet phase is another possibility. This second scenario is reasonable and gains support from our earlier observation of *α*-helix rich conformational compactness of phase-separated papain inside the droplet phase. This conformational compactness of phase-separated papain possibly results in the inactivation of the protease activity. To understand this contrasting and intriguing phenomenon of the protease activity of papain droplets at the microscopic level, we next utilized natural protein substrates.

### Protease Activity toward Natural Serum Albumin Substrates

To decipher the hidden mechanistic aspect of the contrasting protease activities of free papain and papain droplets, we performed a series of spectroscopic and microscopic measurements, including biochemical assays using two natural protein substrates, namely, bovine serum albumin (BSA) and human serum albumin (HSA) (Figure 4a).

**Figure 4.**
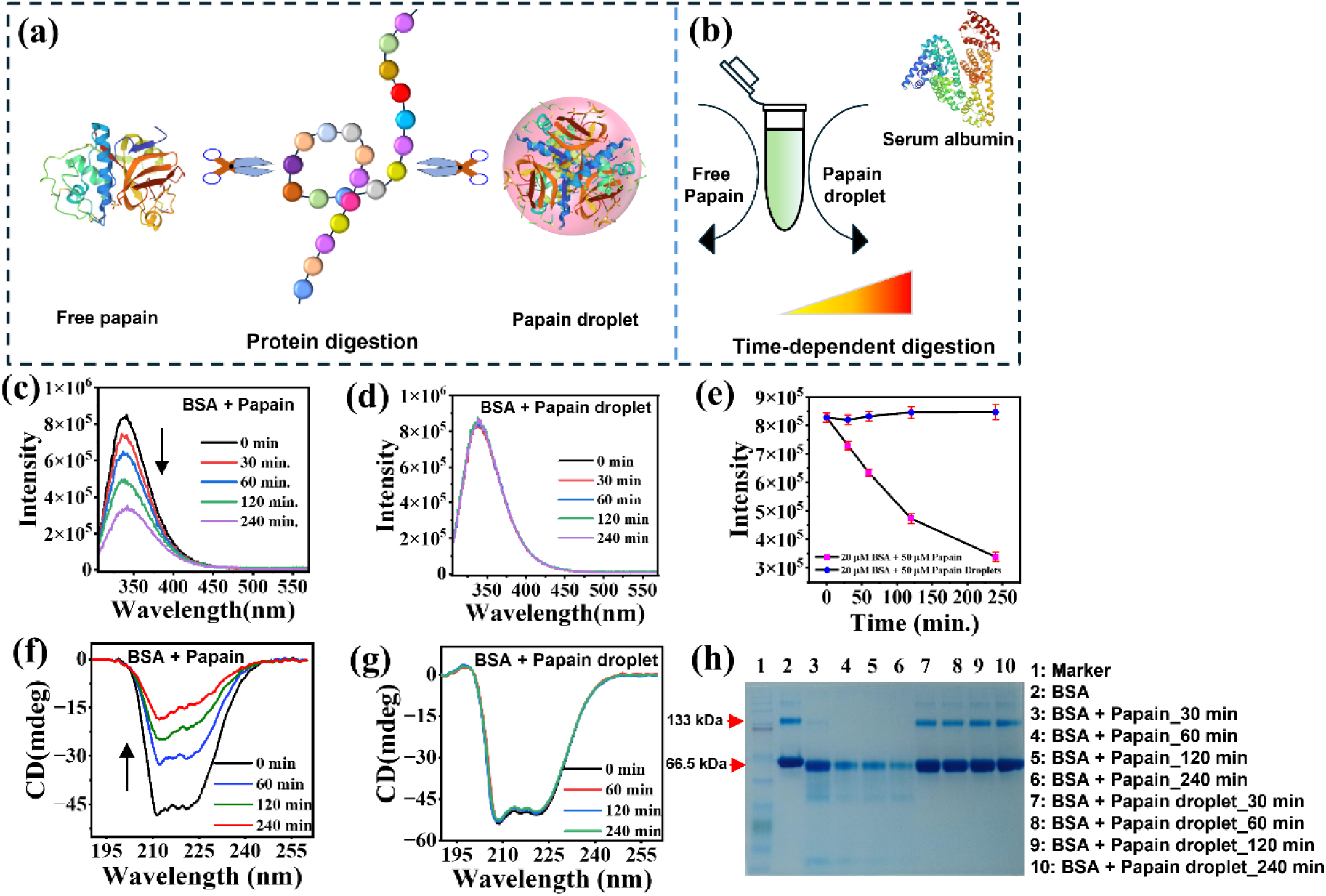
Schematics showing the (a) protease action of free papain and papain droplet on protein substrate, along with (b) time dependent digestion of serum albumin in the presence of free papain and papain droplet. Fluorescence spectra (λ_ex_= 295 nm) of 20 µM BSA as a function of equilibration time in the presence of (c) 8 µM free papain, and (d) papain droplet. (e) Plots of fluorescence intensity of BSA at 340 nm against equilibration time with free papain and papain droplet. Corresponding changes in the far-UV CD spectra of BSA in the presence of (f) free papain and (g) papain droplet over a period of 240 min. (h) Native PAGE data of BSA digestion by free papain and papain droplet. Samples were prepared in 50 mM phosphate citrate buffer (pH 6.0) at 37 °C.

Serum albumins are the most abundant plasma proteins in the blood plasma, and the structure and physicochemical properties of both BSA and HSA have been well explored in the literature.^59–61^ The aqueous solutions of BSA and HSA were equilibrated in the presence of either free papain or papain droplets and the time-dependent digestion was monitored using fluorescence, far-UV CD, and native polyacrylamide gel electrophoresis (PAGE) measurements (Figure 4b). BSA showed an intrinsic fluorescence band centred at 340 nm upon excitation at 295 nm due to the presence of two tryptophan residues (Trp-134 and Trp-213). In the presence of 8 µM papain, the intrinsic fluorescence of BSA gets quenched gradually in a time-dependent manner over a period of 240 min (Figure 4c,e). Moreover, a µnm red shift was observed in the emission spectrum of BSA upon 240 min equilibration with papain. It should be noted that the fluorescence intensity and emission maximum of tryptophan residues in serum albumins are highly sensitive to any conformational fluctuation of the native protein structure. Upon denaturation, tryptophan in serum albumins exhibits quenching in fluorescence with a red-shift in the emission maximum.^60,62^ The observed fluorescence quenching, along with a red shift in the emission maximum of BSA suggests time-dependent denaturation of BSA by papain. In contrast, no noticeable changes in the fluorescence spectra of BSA were observed in the presence of papain droplets under similar experimental conditions (Figure 4d,e). Similar findings were also observed for HSA in the presence of free papain and papain droplets (Figures S13 and S14). This contrasting observation is similar to those observed in the case of the synthetic substrate BAPNA. To gain further insight into the time-dependent digestion, we recorded the far-UV CD spectra of BSA and HSA in the presence of either free papain or papain droplets. The CD spectrum of BSA is characterized by two minima at 208 and 222 nm, signifying an *α*-helix rich globular structure (Figure 4f). Notably, the ellipticity decreases gradually in the presence of papain over a period of 240 min, signifying a decrease in the *α*-helix content as reported previously.^40^ In contrast, the far-UV CD spectra of BSA in the presence of papain droplet remain unaltered as a function of equilibration time. Similar CD spectral changes were observed for HSA in the presence of free papain and papain droplets (Figures S15 and S16). Taken together, these spectral changes suggest that free papain is effective at perturbing the native conformations of serum albumins; however, phase-separated papain is not able to alter the native structures of serum albumins. To know whether these spectral changes are due to progressive digestion of serum albumins by papain, we performed native PAGE. Figure 4h shows the native PAGE of BSA in the absence and presence of free papain and papain droplets at different time intervals. In the absence of papain, BSA showed an intense band at 66.5 kDa and a weak band at 133 kDa due to the presence of monomers and dimers, respectively (Figure 4h; lane: 2).^61^ Very weak bands at higher molecular weights indicate the presence of higher order oligomers of BSA. Similar bands were also observed for HSA in native PAGE (Figure S14; lane: 2). Notably, equilibration of BSA samples with 8 µM papain for different time intervals resulted in appreciable changes in the band intensities of both the monomeric and oligomeric BSA (Figure 4h; lanes: 3–6). Upon 240 min of equilibration, we observed the complete disappearance of the oligomeric bands along with a significant reduction in the monomeric band intensity. These changes were accompanied by the simultaneous appearance of lower molecular weight bands. Similar changes were also observed for HSA in the presence of free papain (Figure S17; lanes: 2–5). These observations authenticate time-dependent digestion of native serum albumins by free papain over a period of 240 min. In contrast, the band intensities of both serum albumins were found to remain unaltered in the presence of papain droplets over a period of 240 min under similar experimental conditions (Figures 4h; lanes: 7–10 and S17; lanes: 6–9). Moreover, no new bands at lower molecular weight were observed upon equilibration with papain droplets even for a period of 240 min. These findings substantiate the lack of any protease activity of phase-separated papain toward serum albumins.

Next, to directly visualize this contrasting protease activity of free papain and papain droplets, we microscopically monitored the protease digestion and fragmentation of FITC-labeled serum albumins by RBITC-labeled papain and papain droplets using CLSM. We equilibrated 20 µM FITC-labeled BSA with 8 µM RBITC-labeled papain or papain droplets at different time intervals, and subsequently, CLSM images were captured (Figure 5a). Upon 1 and 3 h of digestion, we mainly observed diffused fluorescence signals in both red and green channels (Figure 5b). However, beyond 6 h of digestion, random aggregated structures in both red and green channels were observed up to day 1 (d 1). The merged images revealed the presence of both BSA and papain in these aggregated structures. Similar CLSM images were also observed for HSA in the presence of free papain (Figure S18). The diffused fluorescence signals observed till 3 h of digestion mainly originate from the undigested intact proteins along with the peptide fragments. Over time, these peptide fragments undergo random aggregation to yield aggregated structures. At pH 6.0, the opposite charges of papain (pI ∼8.75) and BSA (pI ∼ 4.7) fragments may facilitate the observed random aggregation. In contrast, a drastically different scenario was observed for RBITC-labeled papain droplets (Figure 5c).

**Figure 5.**
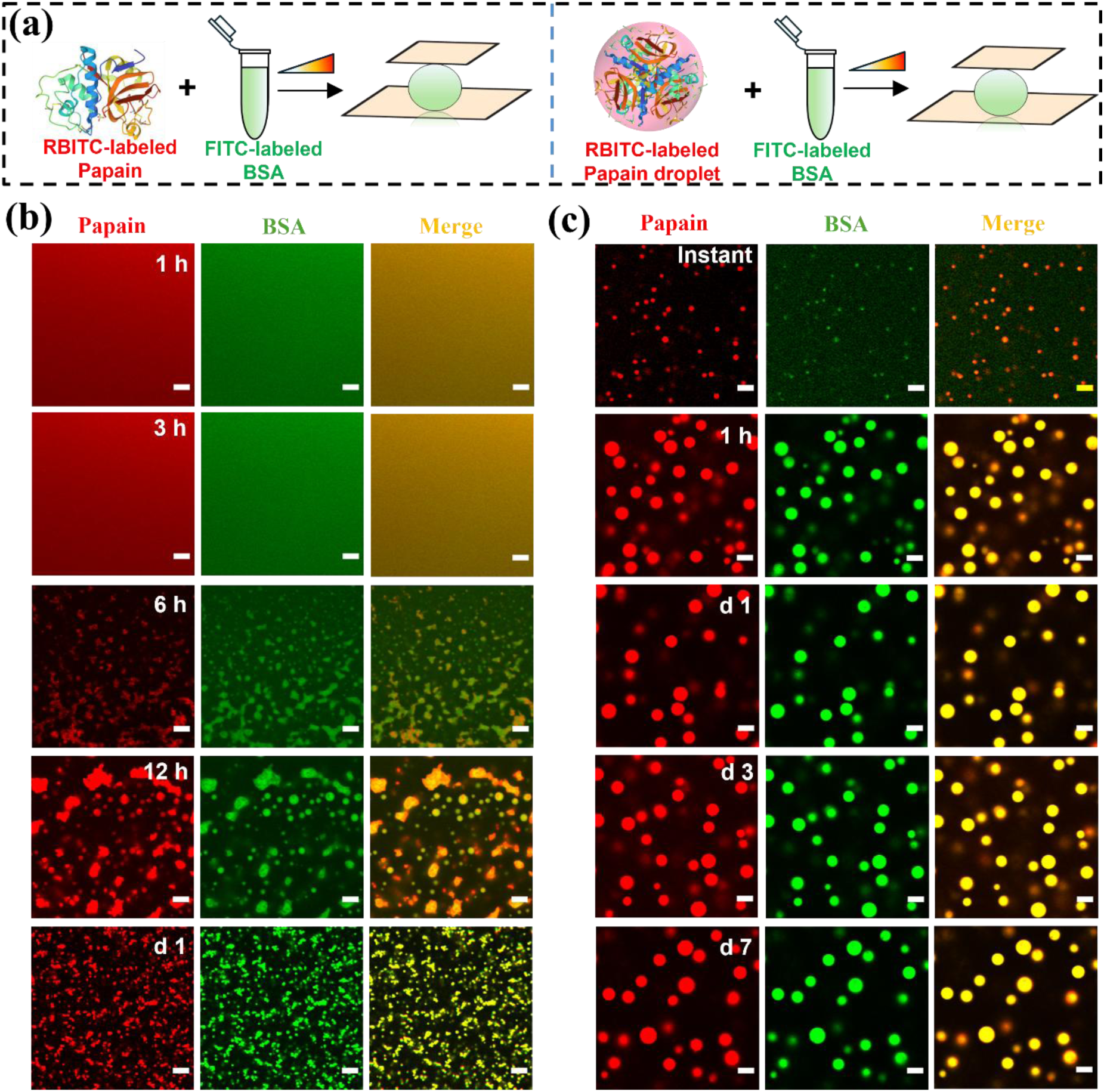
(a) Schematics showing the protease assays performed under CLSM with FITC-labeled BSA in the presence of either RBITC-labeled free papain or papain droplet. CLSM images showing the time dependent digestion of 20 µM FITC-labeled BSA by 8 µM RBITC-labeled (b) free papain and (c) papain droplets. Samples were prepared in 50 mM phosphate-citrate buffer (pH 6.0) at 37 °C. Scale bars correspond to 5 µm.

When CLSM images were captured instantly upon mixing pre-formed RBITC-labeled papain droplets with FITC-labeled BSA, we observed red emitting, uniform spherical papain droplets along with a diffused green signal from FITC-labeled BSA. In addition, a few weak green emissive fluorescence puncta were also observed in the green channel. These green puncta possibly originate due to the partitioning of BSA into the papain droplets, as revealed by the colocalization of red and green emission in the merge image. Upon 1 h of equilibration, we observed complete partitioning of BSA into the papain droplets, as evident from the uniform yellow signals from the droplet interior in the merge image. Interestingly, the fluorescence signals of both papain and BSA remained intact even for a period of 7 days (Figure 5c), suggesting a lack of any time-dependent digestion of BSA by phase-separated papain. These intriguing findings suggest that even though BSA is spontaneously partitioned inside the papain droplets due to their opposite charges, phase-separated papain is not able to digest BSA. Similar results were also observed for HSA when equilibrated with pre-formed papain droplets (Figure S19). This contrasting behavior of papain upon phase separation could be due to the conformational alteration-induced inactivation of its active Cys-25 residue via intermolecular disulfide linkage. To authenticate this argument, we performed a range of control assays using various spectroscopic and microscopic techniques.

### L-cysteine Treatment Activates Papain Droplets

To authenticate the conformational alteration-induced disulfide linkage formation in papain droplets, we performed phase separation and protease assays in the presence of L-cysteine (L-Cys), a well-known natural disulfide linkage disruptor.^20,63^ The catalytically active form of papain contains a sulfhydryl moiety (Cys-25), which acts as a nucleophile to attack the carbonyl carbon of the amide bond (Figure 6a). However, it is known that cysteine proteases can effectively transform into an inactive form via the formation of a disulfide linkage involving the active Cys-25 residue (Figure 6a).

**Figure 6.**
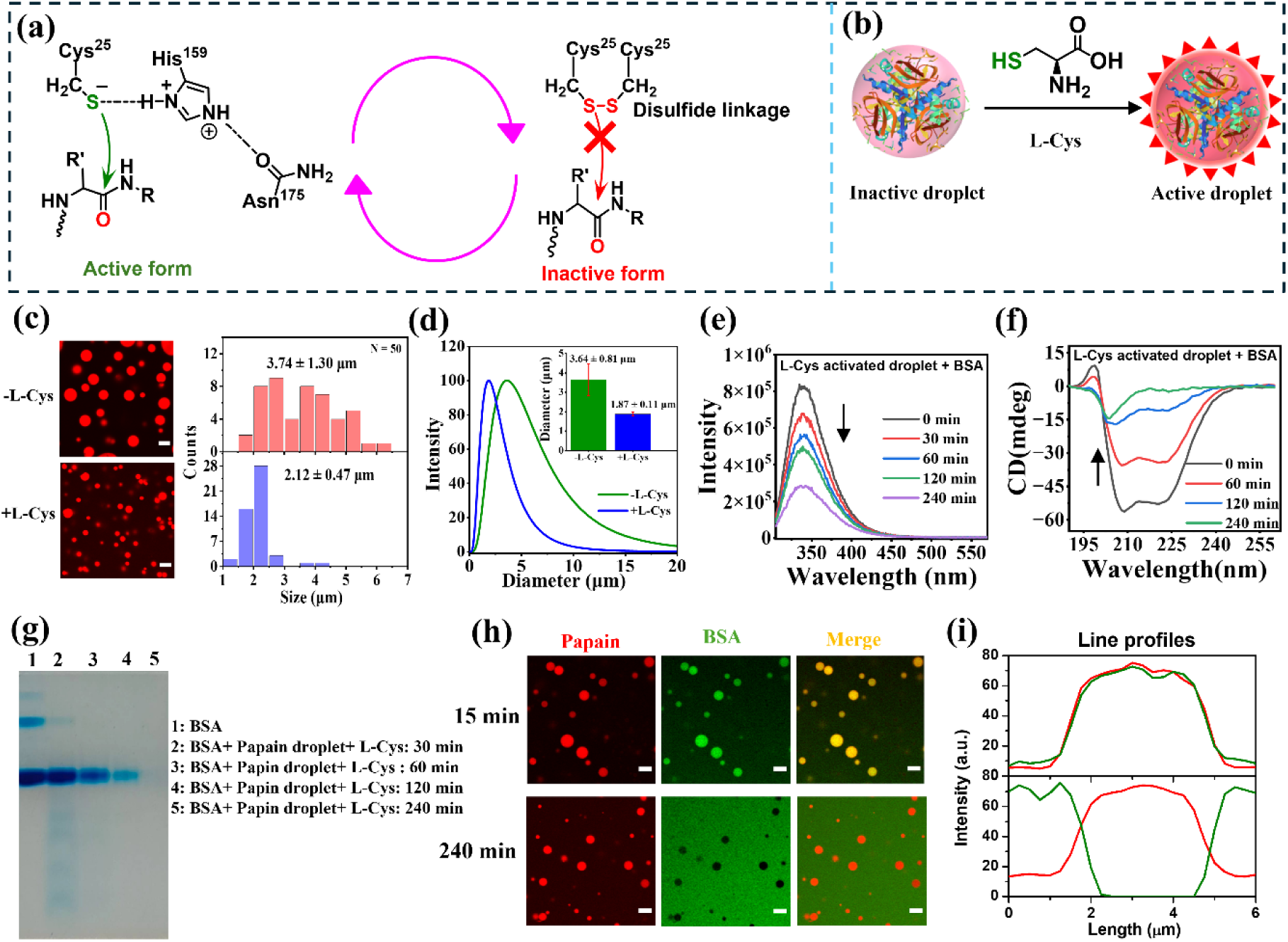
Schematics showing the (a) interconversion of active to inactive form and vice versa involving the Cys-25 residue of papain. (b) The effect of L-Cys treatment on the inactive papain droplet. (c) Confocal images and corresponding size distributions of 8 µM RBITC-labeled papain droplets in the absence and presence of 5 mM L-Cys. (d) DLS size distribution profiles of papain droplets in the absence and presence of 5 mM L-Cys. Time dependent digestion of 20 µM BSA by 8 µM activated papain droplet as revealed from (e) fluorescence spectra (*λ*_ex_ = 295 nm), (f) far-UV CD spectra, and (g) Native PAGE. (h) CLSM images showing the digestion of 20 µM BSA by 8 µM activated papain droplet at time intervals of 15 and 240 min. (i) Corresponding intensity line profiles of activated papain droplets at 15 and 240 min estimated from the green and red channels. Scale bars correspond to 5 µm.

It is highly possible that upon phase separation, conformationally altered papain forms a disulfide linkage inside the droplet phase. To know the molecular origin of this inactivation, we first utilized L-Cys to understand its impact on the LLPS of papain (Figure 6b). Papain droplets were treated with different concentrations of L-Cys and equilibrated for 10 min before any measurements. Although L-Cys does not inhibit the phase separation of papain, the effective size of papain droplets decreased noticeably in the presence of 5 mM L-Cys (Figure 6c). While the estimated mean size of papain droplets was found to be 3.74 ± 1.30 µm in the absence of L-Cys, it decreased to a value of 2.12 ± 0.47 µm in the presence of 5 mM L-Cys (Figure 6c). Notably, the mean droplet size saturated beyond 5 mM L-Cys and droplets remained intact even at 20 mM L-Cys (Figure S20), signifying that LLPS of papain is not driven by disulfide linkage, but rather the effect is associated with the post LLPS process. It should be noted that the addition of other reducing agents, namely, 5 mM dithiothreitol (DTT) and 5 mM *β*-mercaptoethanol (BME), showed a similar reduction in the mean size of droplets (Figures S21 and S22). The hydrodynamic size estimated from DLS also reflected a similar reduction in the mean droplet size from 3.64 ± 0.81 (PDI: 0.509) to 1.87 ± 0.11 µm (PDI: 0.345) upon addition of 5 mM L-Cys (Figure 6d). Next, we evaluated the protease activity of L-Cys-treated papain droplets using serum albumins as natural substrates. Interestingly, L-Cys-treated droplets were found to significantly quench the intrinsic fluorescence of serum albumins over a period of 240 min (Figures 6e and S23). This fluorescence quenching was accompanied by appreciable conformational alterations of serum albumins as evident from the decrease in ellipticity at 208 and 222 nm (Figures 6f and S24). Native PAGE was performed to authenticate the time-dependent digestion of serum albumins by L-Cys-treated papain droplets. Interestingly, L-Cys-treated papain droplets were found to be remarkably efficient in digesting the serum albumins within a period of 240 min (Figures 6g and S25). Both the monomeric and oligomeric bands of serum albumins completely disappeared upon equilibration with L-Cys-treated papain droplets for a period of 240 min. This is in sharp contrast to those observed for untreated papain droplets, suggesting that L-Cys treatment transforms inactive droplets into active droplets. More importantly, the protease activity of activated papain droplets was found to be much more efficient than that of free papain, as revealed by the native PAGE band intensity after 240 min of digestion. To gain a microscopic picture of this highly efficient protease pathway, we performed CLSM imaging using FITC-labeled serum albumins. Papain droplets were activated with 5 mM L-Cys and equilibrated for 10 min before adding serum albumins. Initially, both BSA and HSA partitioned into the RBITC-labeled L-Cys-activated papain droplets within 15 min of mixing, as revealed from CLSM images (Figures 6h and S26). Within 240 min of mixing, we noticed intact red emitting papain droplets in a diffused green background (Figures 6h and S26). The diffused green background suggests efficient digestion and fragmentation of FITC-labeled serum albumins within the activated papain droplets. The resulting smaller fragments emerged from the droplet phase and were distributed uniformly as soluble components. The intensity line profiles at the initial and later stages of BSA digestion by activated papain droplets further highlight the efficient fragmentation within the droplet phase (Figure 6i).

Based on these findings, we propose a mechanistic picture of the LLPS-mediated protease activity of papain and its regulation by L-Cys (Scheme 1). Upon macromolecular crowding, papain undergoes spontaneous LLPS via enthalpically driven homotypic hydrophobic protein-protein interactions to yield phase-separated droplets.

**Scheme S1.**
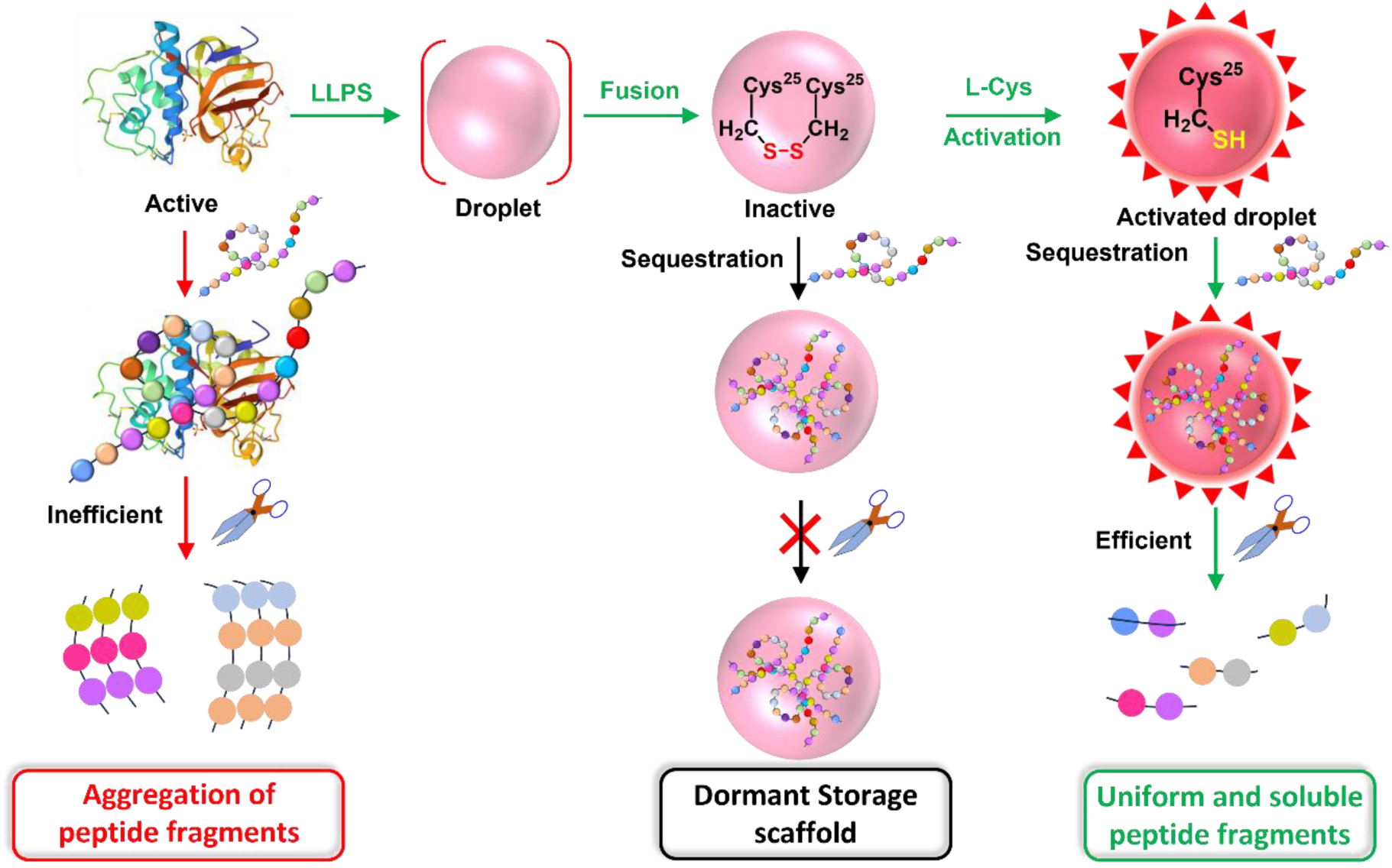
Illustration of Spatio-Temporal Regulation of the Protease Activity of Papain via LLPS and L-Cys-mediated Activation.

Upon phase separation, papain acquires a more compact *α*-helix rich conformational state inside the droplet phase. These droplets are found to be stable over a broad range of pH (3.0–11.0) and temperature (4–90 °C), indicating their robust microstructure. Upon aging, papain droplets undergo spontaneous fusion to yield larger-sized droplets. Although free papain is found to be active toward protease activity with both synthetic and natural substrates, digestion is inefficient and leads to irregular fragmentation and random aggregation of protein fragments. On the other hand, papain droplets are found to be inactive towards protease activity and remain dormant towards both synthetic and natural protein substrates. Our findings indicate that this inactivation is due to the disulfide linkage of phase-separated papain, which is a consequence of fusion mediated intermolecular protein-protein interactions inside the droplet phase. Upon activation with L-Cys, papain droplets exhibit enhanced protease activity and surpass the activity of free papain under similar experimental conditions. In addition, digestion of serum albumins with activated papain droplets leads to the release of uniform and soluble smaller fragments. This highly efficient activity inside the activated papain droplets could be due to the confinement-induced enhanced enzyme-substrate interactions as observed previously in other synthetic reactors. To the best of our knowledge, the present study is the first to report the hidden role of L-Cys-mediated regulatory pathway during the protease activity of papain.

## SUMMARY

In the present study, we have discovered a new regulatory pathway for the protease activity of papain under cell mimicking heterogeneous and crowded environments. We show that the spontaneous LLPS of papain under macromolecular crowding proceeds via enthalpically driven hydrophobic protein-protein interactions in a broad range of pH and temperature. Phase-separated papain inside the droplet phase attains a more compact *α*-helix rich conformation as revealed from the secondary structure analyses. While free native papain is found to be active toward synthetic and natural protein substrates, the phase-separated papain fails to exhibit any protease activity. Our findings reveal that this contrasting behavior is due to the fusion-mediated intermolecular disulfide linkage formation involving active Cys-25 residues of phase-separated papain. Moreover, we show that these dormant droplets can be reactivated in the presence of L-cys. These activated droplets exhibit enhanced efficacy toward protease activity to yield uniform soluble peptide fragments, which is superior to that observed for free native papain. These findings provide a new innovative platform for technological advancement in the food industry.

## EXPERIMENTAL SECTION

### Materials

Human serum albumin (HSA), bovine serum albumin (BSA), polyethylene glycol (PEG 8000), mPEG-NH_2_ (MW 5000), dextran 70, Ficoll 400, rhodamine B isothiocyanate (RBITC), fluorescein-5-isothiocyanate isomer-I (FITC), Hellmanex III, *N_α_*-benzoyl-L-arginine *p*-nitroanilide hydrochloride (BAPNA), 1,6-hexanediol and adenosine 5’-triphosphate disodium salt hydrate were purchased from Sigma-Aldrich. Tri-sodium citrate, citric acid anhydrous, sodium dihydrogen phosphate monohydrate (NaH_2_PO_4_·H_2_O), disodium hydrogen phosphate heptahydrate (Na_2_HPO_4_·7H_2_O), sodium carbonate anhydrous, sodium azide (NaN_3_), sodium thiocyanate (NaSCN), ammonium sulfate ((NH_4_)_2_SO_4_), sodium chloride (NaCl), sodium hydroxide (NaOH) and sodium salt of ethylene diamine tetra acetate (EDTA) were purchased from Merck. Papain 5x USP ex. Papaya Latex, 30000 USP U/mg powder, sodium bicarbonate extra pure, and Ficoll 400k were purchased from SRL chemicals. Glycine was purchased from TCI chemicals. L-cysteine was purchased from Loba Chemie. Potassium dichromate and sulfuric acid were purchased from SDFCL. All the chemicals were used without any further purification. Eco Testr pH1 pH meter was used to adjust the final pH (±0.1) of all the buffer solutions. Milli-Q water was obtained from a Millipore water purifier system (Milli-Q integral).

### Characterization Techniques

#### UV-Visible and Fluorescence Spectroscopy

UV-vis absorption spectra were recorded in a quartz cuvette (1 × 1 cm) using a Varian Carry 100 Bio UV–visible spectrophotometer from Agilent Technologies. The fluorescence spectra were recorded in a quartz cuvette (1 cm × 1 cm) using a HORIBA Jobin Yvon, model FM-100 Fluoromax-4 Spectrofluorometer. The slit width was kept at 1 nm.

#### Dynamic Light Scattering (DLS) Measurements

DLS measurements were performed using a Brookhaven particle size analyzer (model NanoBrook Omni) to estimate the hydrodynamic diameters of papain droplets in the absence and presence of L-Cys. Initial buffer solutions were filtered through a 0.22 µm syringe filter (Whatman) before any sample preparation.

#### Fourier-Transform Infrared (FTIR) Spectroscopy

FTIR measurements were performed to determine the secondary structure of papain by using a Bruker spectrometer (Tensor-27). An aliquot of 50 μL from a 1.66 μM stock solution of papain in the absence and presence of 10% PEG 8000 in a pH 6.0 phosphate citrate buffer was used for the FTIR measurement. The spectra were recorded in the range of 4000–400 cm^−1^. The Fourier self-deconvolution (FSD) method was used to deconvolute the spectra corresponding to the wavenumbers 1700–1600 cm^−1^. The Lorentzian curve fitting was done to fit the spectra by using the Origin 2025b software. The experiments were performed thrice with similar observations.

#### Confocal Laser Scanning Microscopy (CLSM)

The confocal images were obtained using an inverted confocal microscope, an Olympus Fluo-view (model FV1200MPE, IX-83), through an oil immersion objective (100×, 1.4 numerical aperture). The samples were excited with two different lasers (488 and 561 nm) by using appropriate dichroic and emission filters in the optical path. Confocal images of the liquid phase experiment were captured by drop-casting a 10 μL aliquot of the sample solution onto a cleaned glass slide and sandwiching it with a Blue Star coverslip. The edges of the coverslips were sealed with a minimal amount of commercially available nail paint.

#### Circular Dichroism (CD) Spectroscopy

CD spectra were recorded on a JASCO J-815 CD spectropolarimeter using a quartz cell with a path length of 1 mm and a scan range of 190–260 nm. Scans were recorded with a slit width of 1 nm and a speed of 50 nm/min. The CD spectrum of papain was recorded at a concentration of 1.66 μM in a pH 6.0 phosphate citrate buffer. For protein digestion assays, the CD spectra of serum albumins were recorded at a concentration of 4 µM of serum albumins in the presence of 1.66 µM of papain.

#### Native PAGE

Equal amounts of protein samples were mixed with 2X gel loading dye (32 mM Tris-HCl, pH 6.8, 15% (v/v) glycerol, 0.01% bromophenol blue). The samples were resolved in 10% native PAGE gel (Stacking 5 mL: 4% acrylamide, 0.125 mM Tris pH 6.8, 0.1% APS, 0.1% TEMED; Resolving 10 mL: 10% acrylamide, 0.39 mM Tris pH 8.8, 0.1% APS, 0.1% TEMED) by running at a constant voltage of 80 V using the Mini Protean cell system (Bio-Rad). Native PAGE gels were stained with Coomassie Brilliant Blue solution (0.1% Coomassie brilliant blue solution, 50% v/v Methanol, 10% glacial acetic acid and 40% double distilled water) for 2 h and then destaining (30% v/v Methanol, 10% glacial acetic acid and 60% double distilled water) was done for 3–4 times, each wash lasting for 1-2 h.

## Methods

### Prediction of LCDs and IDRs of Papain

To predict the presence of low complexity domains (LCDs) and intrinsically disordered regions (IDRs) in the amino acid sequence of papain protease, we used Simple Molecular Architecture Research Tool (SMART) (http://smart.embl-heidelberg.de/)^44^ and IUPred2A (https://iupred2a.elte.hu/),^45^ respectively. The IUPred2A data were plotted using the Origin software 2025b.

### Prediction of the Probability of LLPS

To predict the probability of spontaneous liquid-liquid phase separation (LLPS) of papain, we utilized the FuzDrop algorithm.^64^ The FuzDrop data were plotted using the origin software 2025b.

### Preparation of Buffer and Crowder Solutions

All the buffer solutions were prepared in the presence of 0.02% sodium azide and sterilized by autoclaving to minimize the growth of bacteria. Different buffer solutions with pH values of 2.0 to 11.0 were prepared by using Milli-Q water. The buffer strength was kept constant at 50 mM. Glycine-HCl buffer (pH 2.0 and 3.0), phosphate-citrate buffer (pH 4.0, 5.0 and 6.0), phosphate buffer saline (pH 7.0, 50 mM NaCl), glycine-NaOH buffer (pH 8.0), and carbonate-bicarbonate buffer (pH 9.0, 10.0, and 11.0) were used individually.

Solutions of different crowders such as 10% (w/v) polyethylene glycol (PEG 8000), 10% (w/v) dextran 70, and 12.5% (w/v) Ficoll 400 were prepared from stock solutions of 40% (w/v) PEG 8000, 40% dextran 70, and 50% (w/v) Ficoll 400, respectively. 20 mg/mL bovine serum albumin (BSA) was prepared from the stock solution of 199.5 mg/mL BSA.

### Labelling of Protein and Crowders with Fluorescent Dyes

The concentration of papain was estimated spectrophotometrically at *λ* = 280 nm using the reported extinction coefficient of 57600 M^-1^ cm^-1^.^65^ Papain was labelled with RBITC and FITC dyes according to an earlier reported method.^33,34^ In short, 300 µM papain was mixed with RBITC or FITC in a molar ratio of 1:2 (papain:dye). The mixture was incubated for 4 h at room temperature, followed by 6 h at 4 °C on a magnetic stirrer with constant stirring at 240 rpm. After the completion of the reaction, the unconjugated dyes were removed using a dialysis membrane (molecular weight cut-off 10–12 kDa) against 50 mM pH 7.4 phosphate buffer at 4 °C for 12 h with regular buffer exchange in 2 h intervals. The same procedure was followed for the labelling of bovine serum albumin using FITC and RBITC dyes.

### Cleaning of Coverslips and Glass Slides for Microscopy

The coverslips and glass slide were first cleaned with 10% chromic acid solution (sulphuric acid and potassium dichromate at 1:1) for 30 min, then cleaned with 2% Hellmanex III solution. Each of these cleaning steps was followed by repeated washing with Milli-Q water. Finally, these washed coverslips and glass slides were rinsed with methanol and dried in a vacuum oven.

### LLPS Assays of Papain in the Presence of Crowders

LLPS assays of papain were performed in the presence of different crowders (PEG 8000, dextran 70, Ficoll 400, and BSA) as a function of papain concentration (100 pM to 50 µM), crowder concentration (1–20%), and incubation time of 24 h at 37 °C. These samples were prepared in 5 mL glass vials and then kept at 37 °C in an incubation chamber with constant temperature regulation.

For the temperature dependent study, samples (1.66 µM papain in the presence of 10% PEG 8000) were prepared in pH 6.0 phosphate-citrate buffer at different temperatures of 4, 25, 37, 50, 70, 90, and 100 °C and incubated for 1 h. Similarly, a pH dependent study was carried out using 10 µM papain in the presence of 10% PEG 8000 in different buffers with pH values of 2.0, 3.0, 4.0, 5.0, 6.0, 7.0, 8.0, 9.0, 10.0, and 11.0 at 37 °C.

The salt-dependent study was performed by adding different concentrations of NaCl (0–3 M), NaSCN (0–3 M), 1,6-hexanediol (0–10%), ATP (0–50 mM), and ammonium sulphate (0–1 M) from their respective stock solutions. Solutions were prepared by dissolving different concentrations of salts/aliphatic alcohol in aqueous phosphate-citrate buffer (pH 6.0) containing 1.66 µM papain and 10% PEG 8000. Solutions were kept at 37 °C for 1 h, and subsequently measurements were performed.

### Protease Activity of Papain

The protease activity of papain and its associated kinetics were monitored by using BAPNA as a synthetic substrate. Activity measurements were performed using 1.66 µM papain in the presence of different concentrations of BAPNA in the concentration range from 50 µM to 10 mM. The UV-vis spectra of BAPNA were recorded at an interval of 2 min for a total period of 10 min. The variation of the absorbance at 405 nm with time was fitted with a linear function, and the slope was estimated. The initial velocity for the hydrolysis of BAPNA by papain was calculated with the value of slope from the linear plot and the reported molar extinction coefficient for *p*-nitroaniline at 405 nm (9960 M^-1^ cm^-1^).^66^ Subsequently, Michaelis-Menten plots were generated by plotting the initial velocity against substrate concentrations. The data were fitted with the Michaelis-Menten equation to estimate the Michaelis constants *K*_m_ and *V*_max_. Finally, the turnover number (*k*_cat_) was calculated using the estimated Vmax and initial enzyme concentration (1.66 µM). For the protease activity measurements using serum albumins as natural protein substrate, activity measurements were performed using 8 µM papain with either 20 µM BSA or HSA in 50 mM phosphate-citrate buffer (pH 6.0) at 37 °C.

## ASSOCIATED CONTENT

### Supporting Information

Confocal images of RBITC-labeled papain droplets as a function of papain concentrations; aging effect of RBITC-labeled papain droplets; intensity line profiles of RBITC-labeled BSA and papain droplets; LLPS as a function of papain and PEG 8000 concentrations; Confocal images of RBITC-labeled papain droplets as a function of equilibration time; deconvoluted FTIR spectra of free papain and papain droplets with PEG 8000, dextran 70, and Ficoll 400; Confocal images of RBITC-labeled papain droplets as a function of pH, temperature and (NH_4_)_2_SO_4_ concentrations; UV-vis spectra of BAPNA in the absence of papain; Fluorescence spectra of HSA in the presence of free papain and papain droplets; far-UV CD spectra of HSA in the presence of free papain and papain droplets; native PAGE of HSA in the absence and presence of free papain and papain droplets; confocal images of time dependent digestion of HSA in the presence of free papain and papain droplets; confocal images of RBITC-labeled papain droplets as a function of L-Cys, DTT, and BME concentrations; fluorescence and CD spectra of HSA in the presence of L-Cys-activated papain droplets; native PAGE of HSA in the absence and presence of L-Cys-activated papain droplets; confocal images of time dependent digestion of HSA in the presence of L-Cys-activated papain droplets (PDF)

## AUTHOR INFORMATION

### Notes

The authors declare no competing financial interests.

## ACKNOWLEDGEMENTS

The authors acknowledge Indian Institute of Technology (IIT) Indore for providing financial support, and infrastructure. This work is financially supported by Council of Scientific and Industrial Research (CSIR) grant no. 01/3108/23/EMR-II and Science and Engineering Research Board (SERB) grant no. CRG/2023/003004. The authors acknowledge SIC, IIT Indore, for instrumental facilities. S.G. acknowledges Department of Science and Technology (DST), India for INSPIRE research fellowship.

