## Supporting Information File for "Biomolecular Condensation and L-Cysteine Signaling Activates Dormant Protease Activity of Papain Droplets"

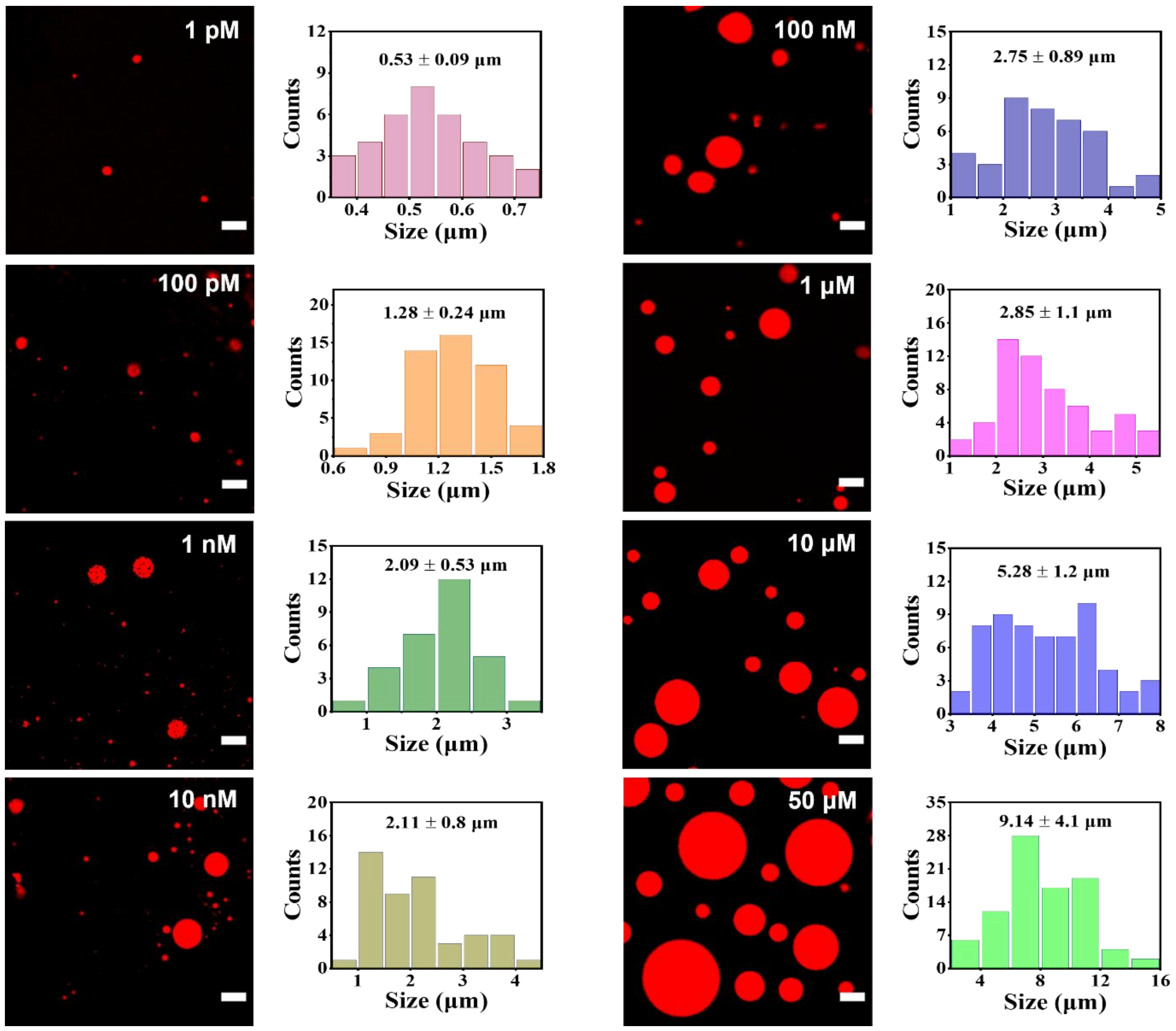


**Figure S1.** Confocal images and associated size distributions of RBITC-labeled papain at different concentrations in the presence of 10% PEG 8000. Samples were prepared in phosphate-citrate buffer at pH 6.0 and incubated at 37 °C for 1 h. Scale bars correspond to 5 µm.

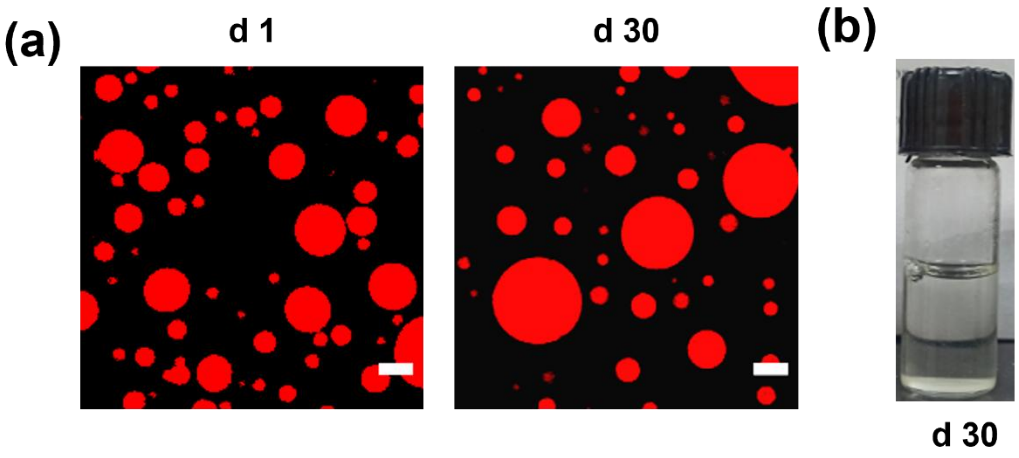


**Figure S2.** (a) Confocal images showing the aging effect of papain droplet over a period of day 30. Scale bars correspond to 5 µm. (b) Daylight photograph of aqueous suspension of papain droplets upon 30 days of aging showing absence of any liquid-to-solid phase transition.

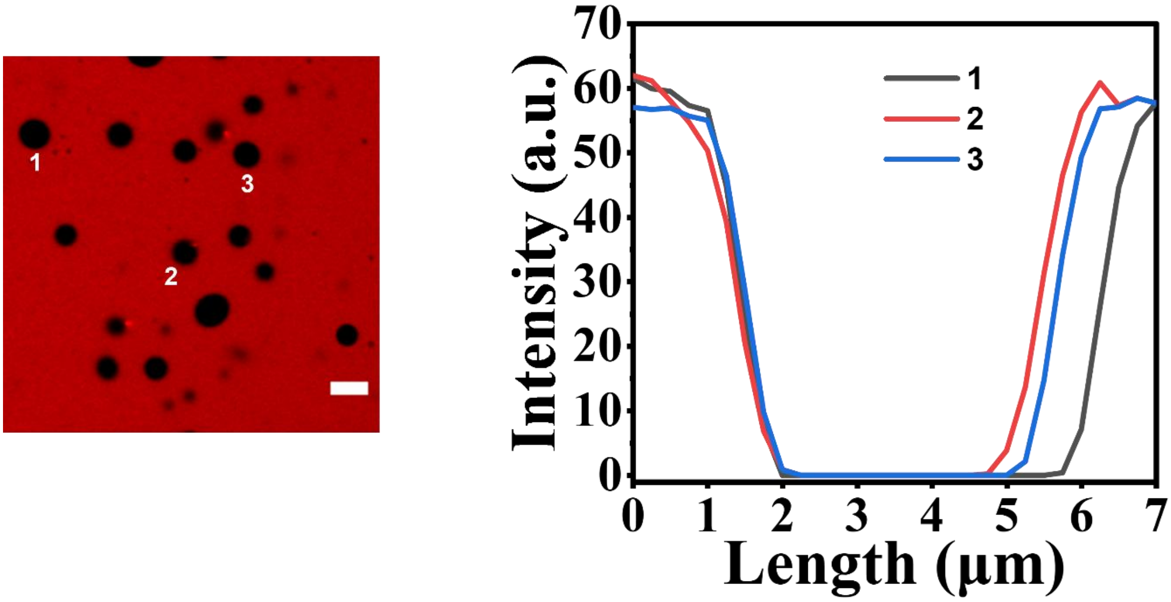


**Figure S3.** Intensity line profiles of unlabeled papain droplets in the presence of RBITC-labeled 10% mPEG-$NH_2$ (MW 5000). Scale bar corresponds to 5 μm.

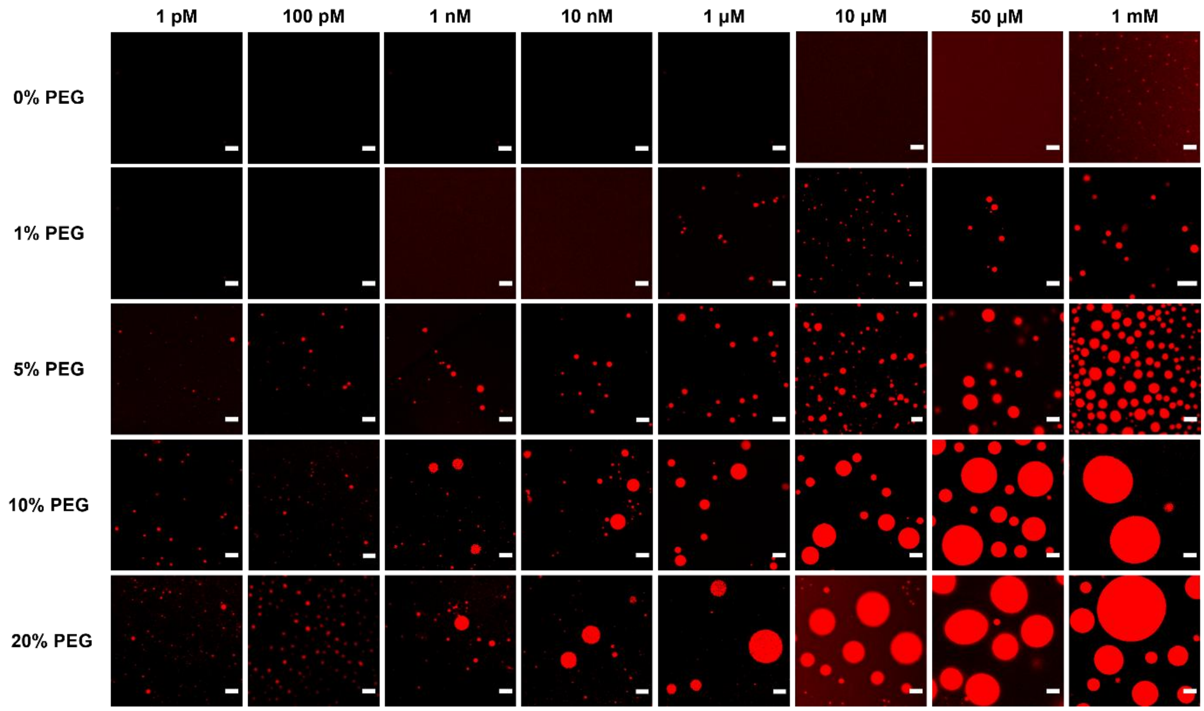


**Figure S4.** Confocal images of RBITC-labeled papain showing the effect of papain and PEG 8000 concentrations on the LLPS. Scale bars correspond to 5 µm.

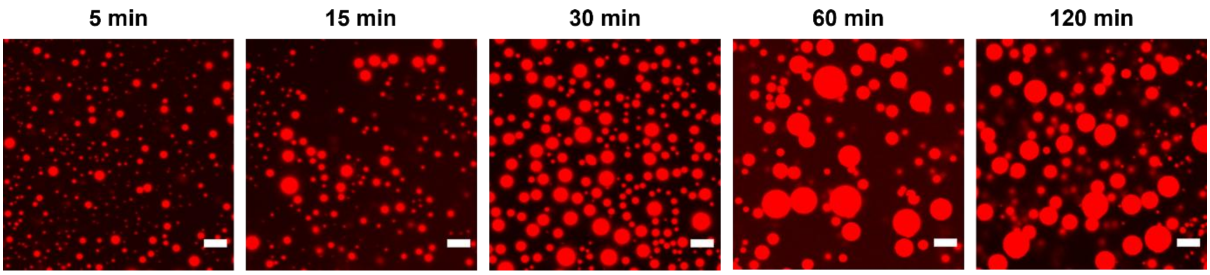


**Figure S5.** Confocal images showing the effect of equilibration time on the size of 1.66 μM papain droplets in the presence of 10% PEG 8000 over a period of 120 min. Scale bars correspond to 5 μm.

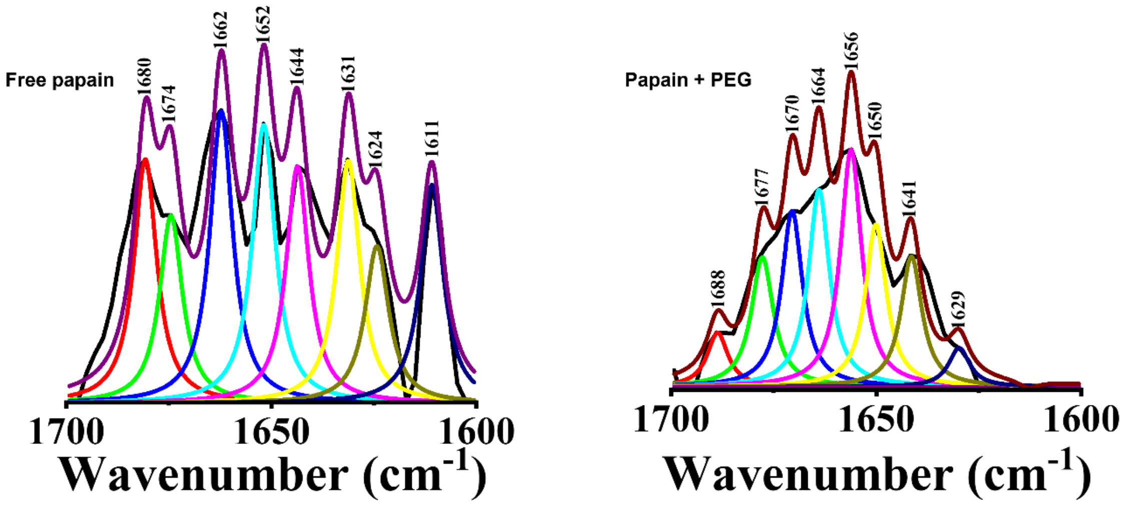


**Figure S6.** Deconvoluted FTIR spectra of 1.66 μM of (a) free papain, and (b) papain droplets with 10% PEG 8000.

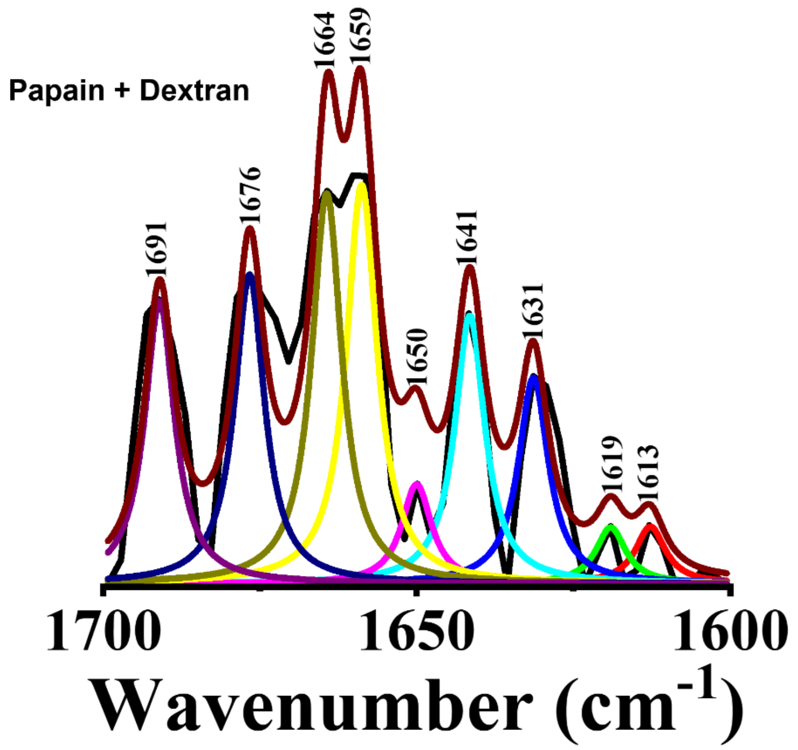


**Figure S7.** Deconvoluted FTIR spectra of 1.66 µM papain with 10% dextran70.

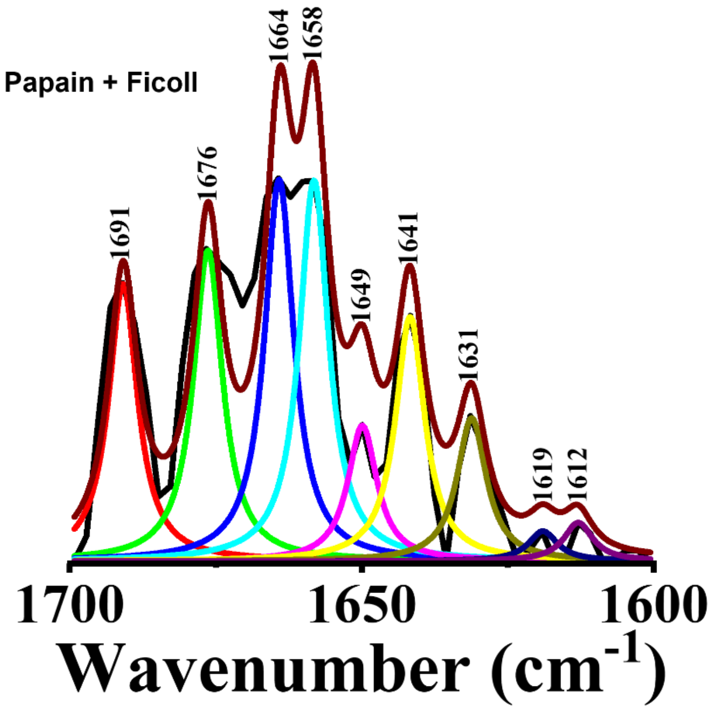


**Figure S8.** Deconvoluted FTIR spectra of 1.66 µM papain with 12.5% Ficoll 400.

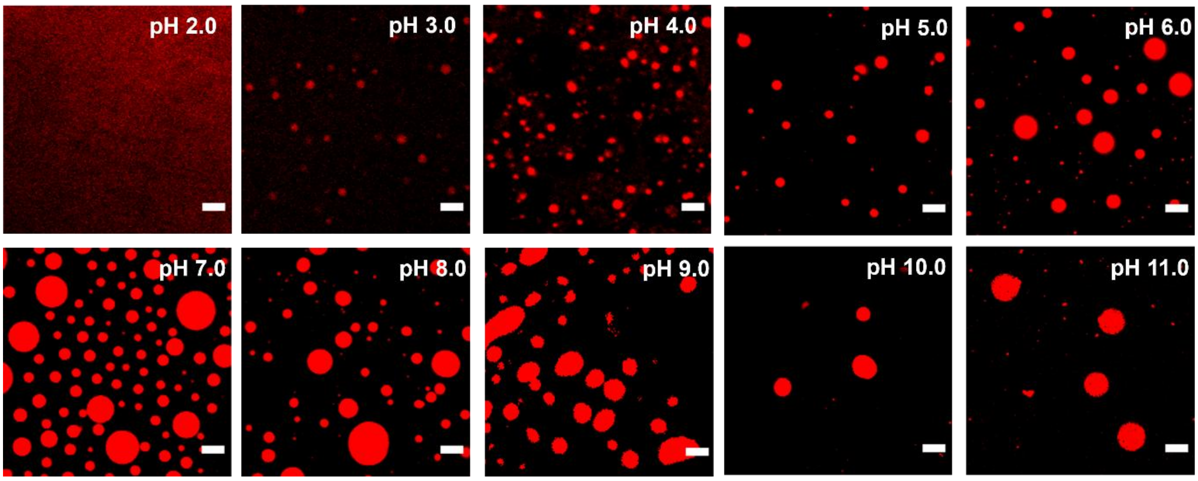


**Figure S9.** Confocal images of RBITC-labeled papain droplets at different pH. Scale bars correspond to 5 μm.

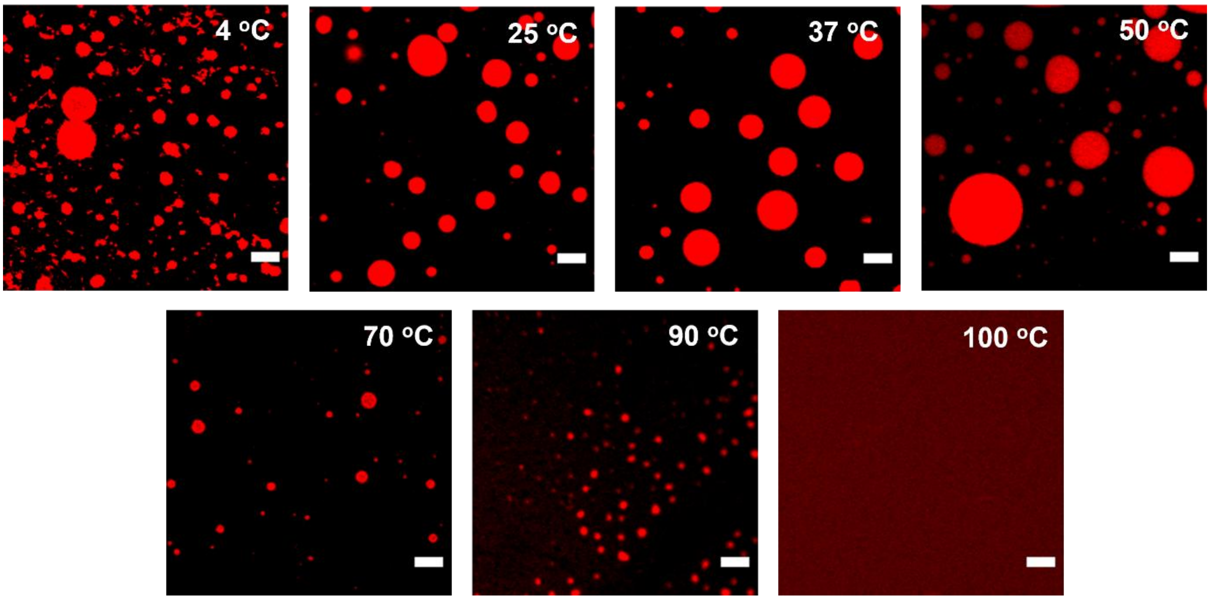


**Figure S10.** Confocal images of papain droplets at different temperatures. Scale bars correspond to 5 μm.

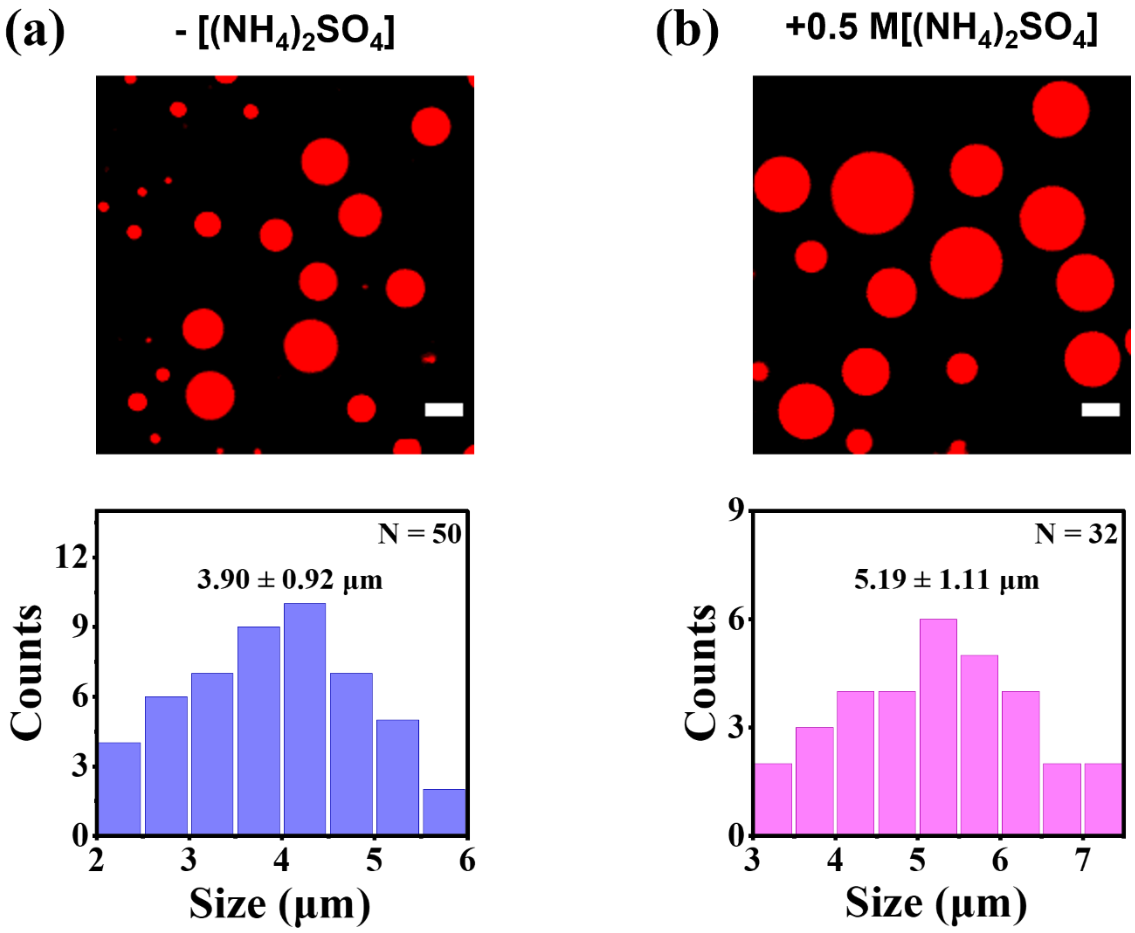


**Figure S11.** Confocal images and associated size distributions of papain droplets in the (a) absence and (b) presence of $(NH_4)_2SO_4$. Scale bars correspond to 5 µm.

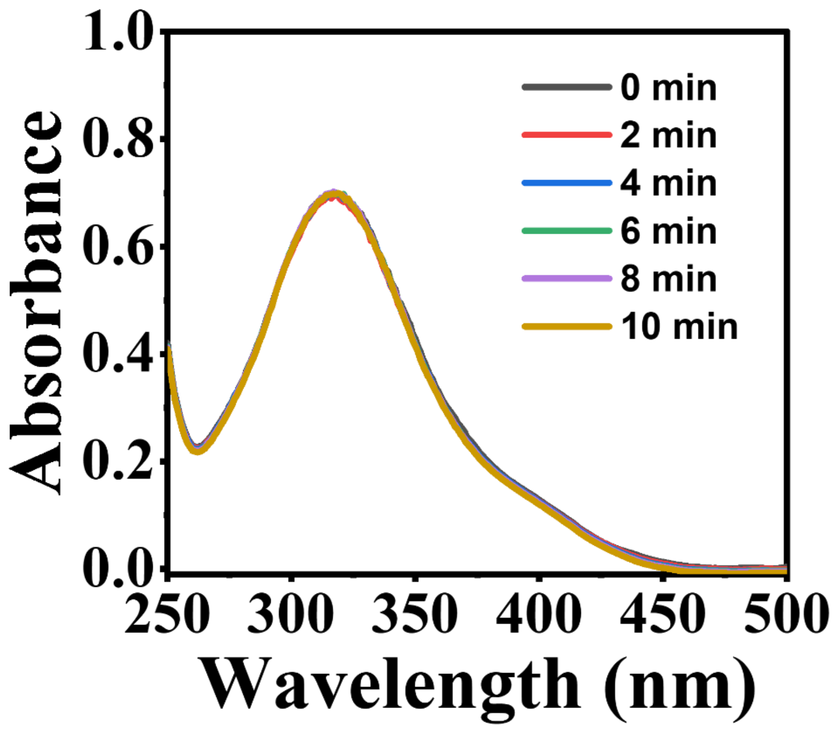


**Figure S12.** Time dependent UV-vis spectra of 50 μM BAPNA in the absence of papain in 50 mM phosphate-citrate buffer (pH 6.0) at 37 $^{o}$C.

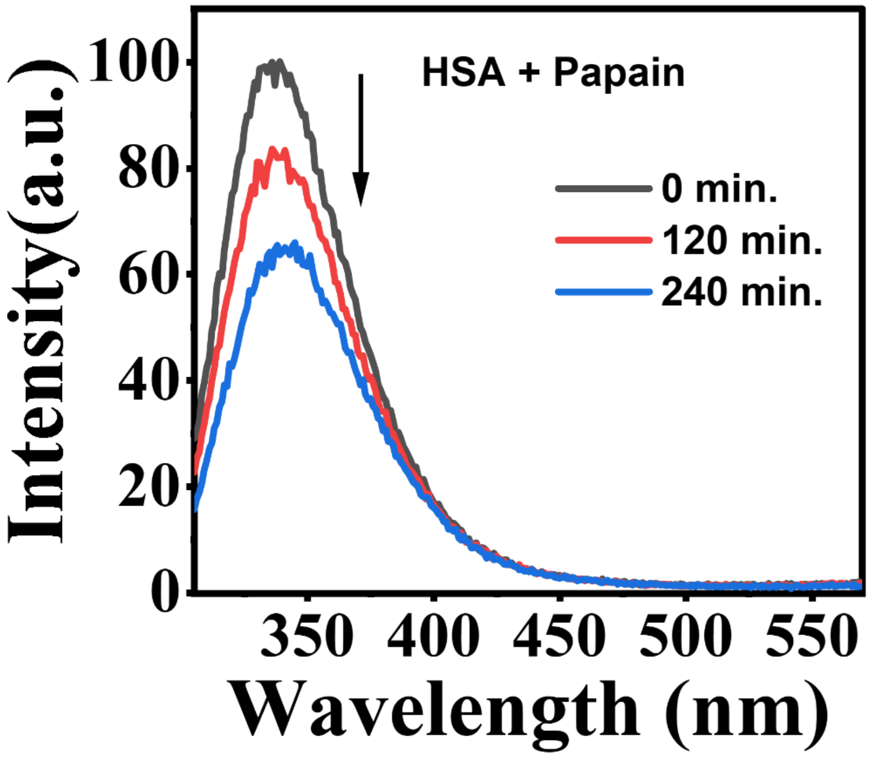


**Figure S13.** Corrected fluorescence spectra ($\lambda_{ex}$ = 295 nm) of 20 μM HSA in the presence of 8 μM papain as a function of digestion time.

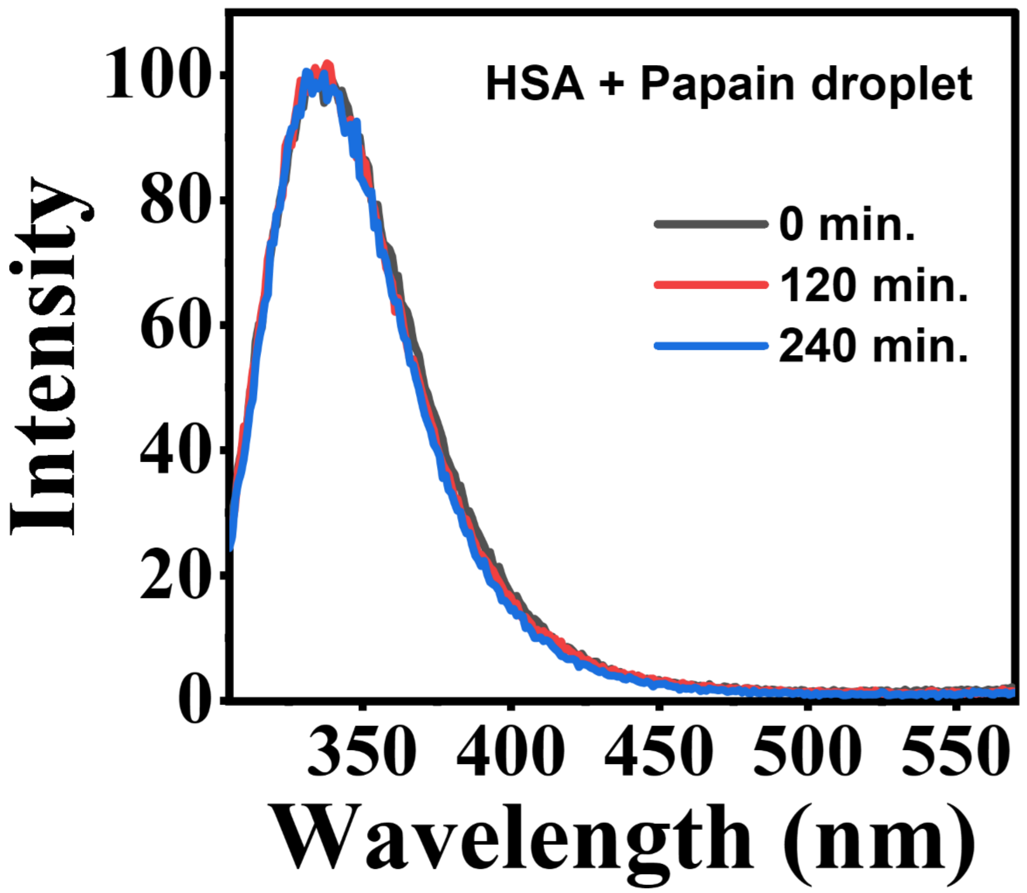


**Figure S14.** Corrected fluorescence spectra ($\lambda_{ex}$ = 295 nm) of 20 µM HSA in the presence of papain droplet (initial concentration of papain was 8 µM) as a function of digestion time.

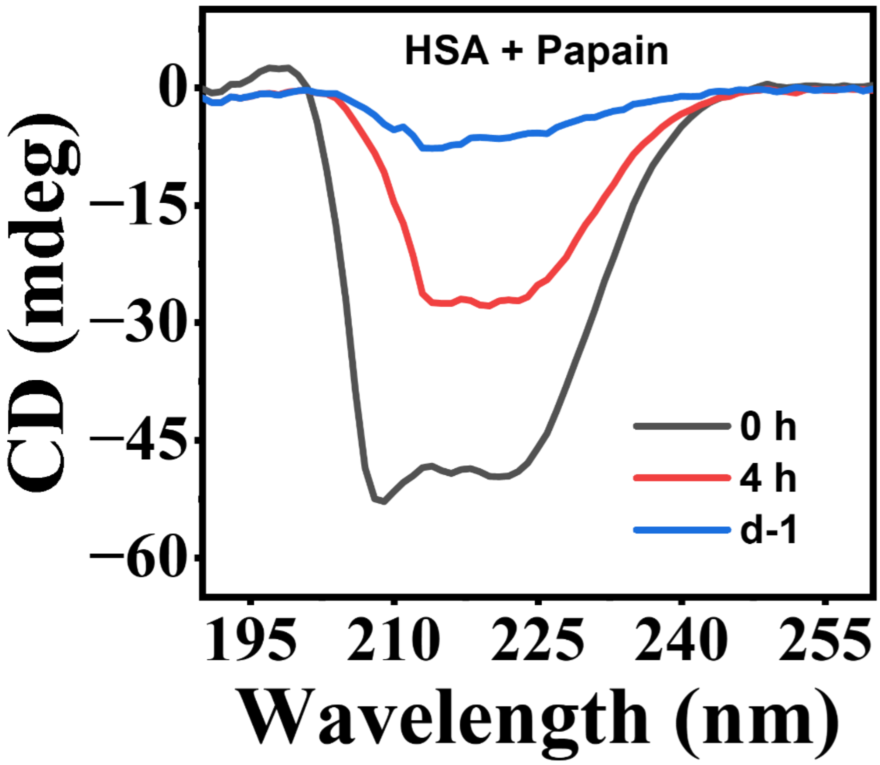


**Figure S15.** Far-UV CD spectra of 4 μM HSA in the presence of 1.66 μM papain.

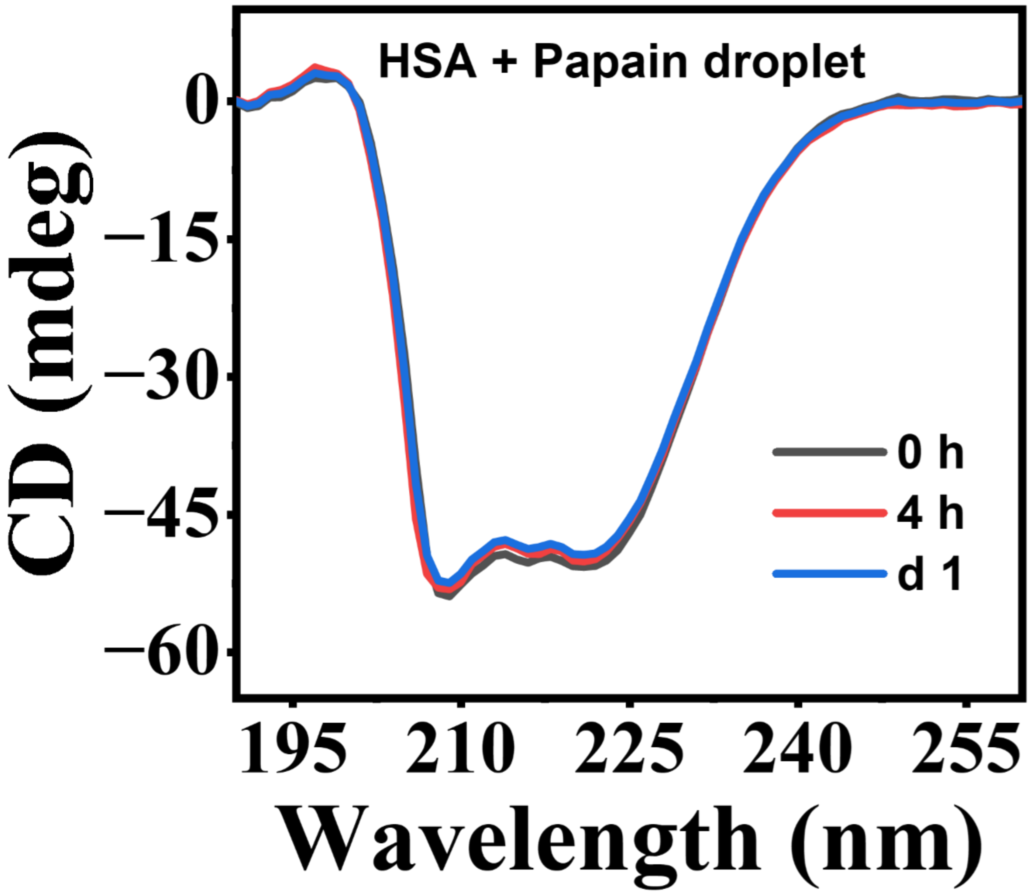


**Figure S16.** Far-UV CD spectra of 4 μM HSA in the presence of papain droplet (initial concentration of papain was 1.66 μM).

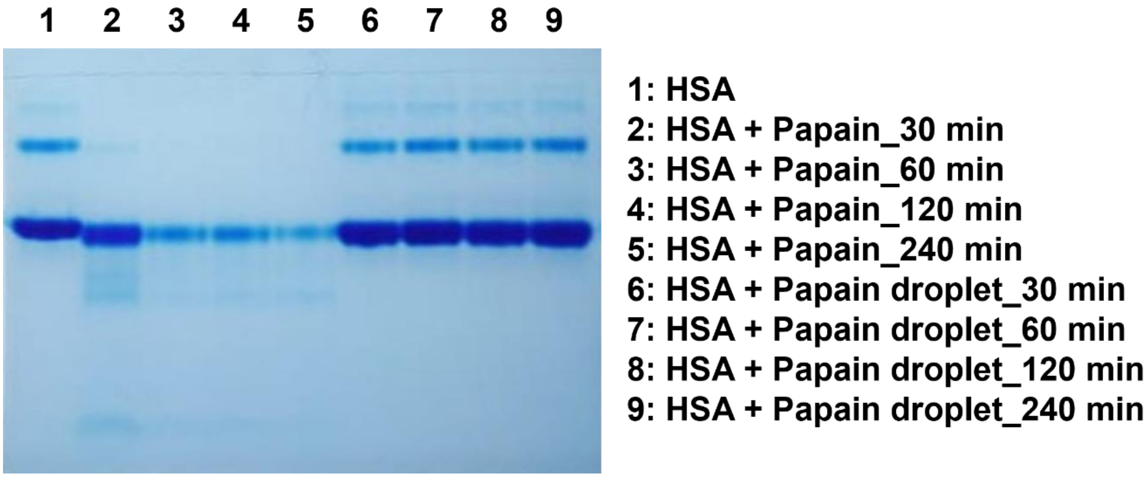


**Figure S17.** Native PAGE data of HSA in the absence and presence of free papain and papain droplets over a period of 240 min.

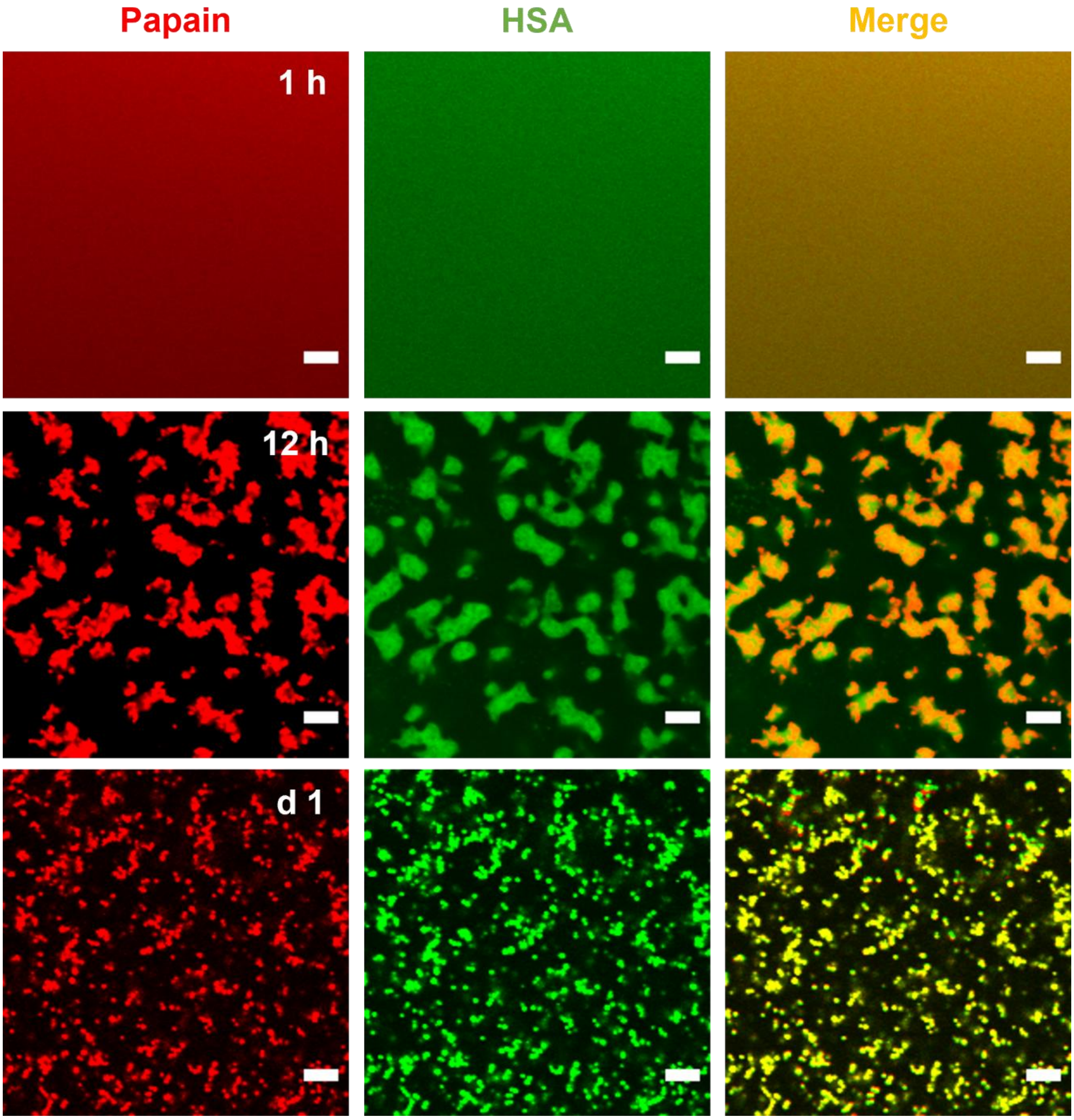


**Figure S18.** Confocal images showing the time dependent digestion of 20 μM FITC-labeled HSA by 8 μM RBITC-labeled free papain. Scale bars correspond to 5 μm.

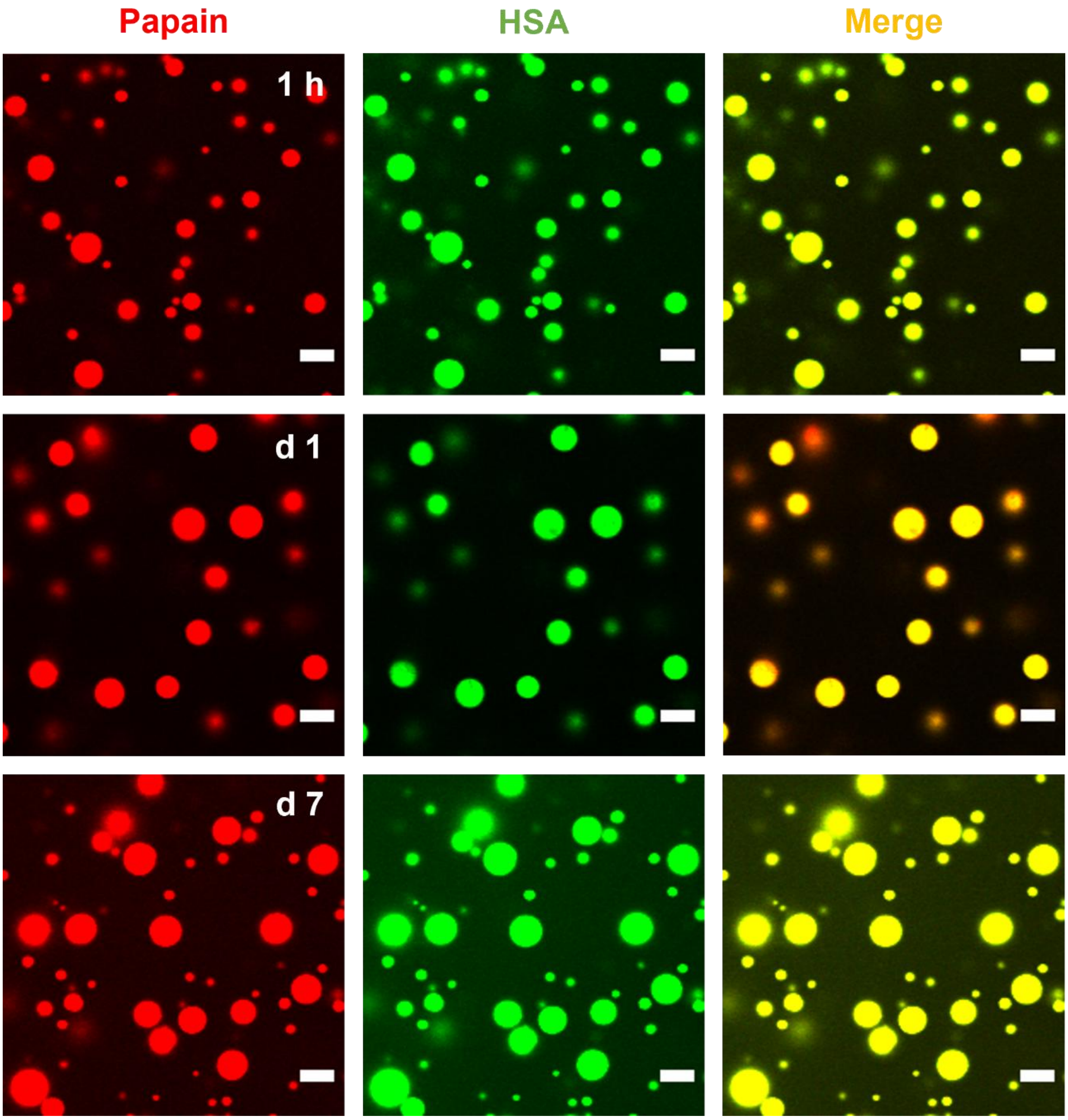


**Figure S19.** Confocal images showing lack of digestion of 20 μM FITC-labeled HSA by RBITC-labeled papain droplets (initial concentration of papain 8 μM) over a period of 7 days. Scale bars correspond to 5 μm.

(a)

(b)

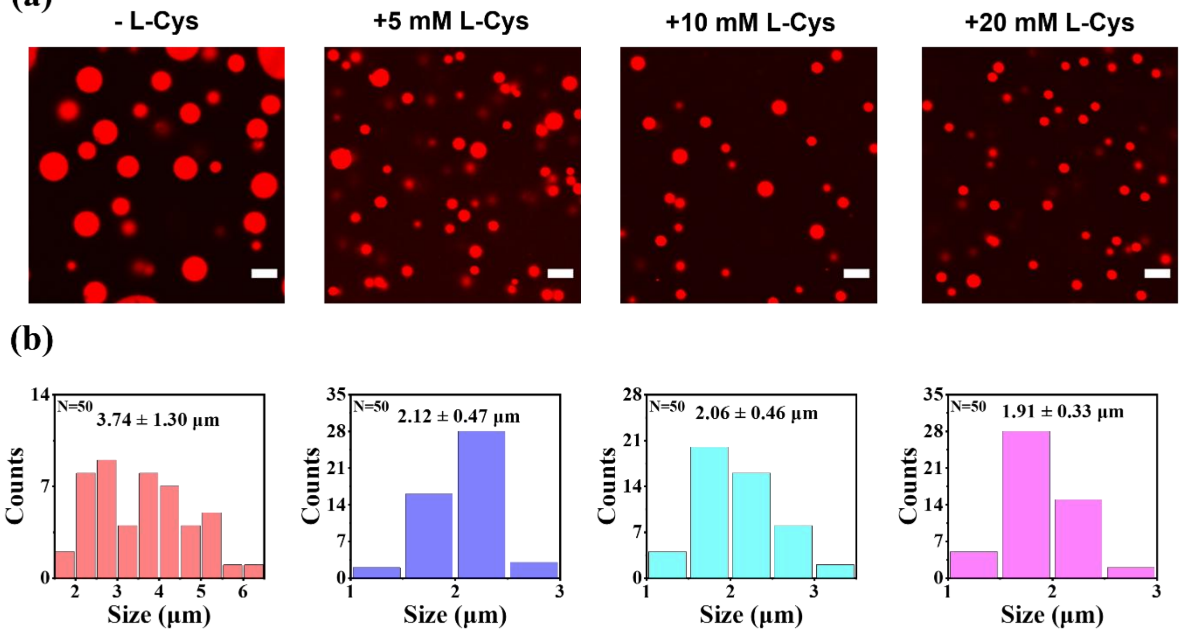


**Figure S20.** (a) Confocal images of RBITC-labeled papain droplets in the absence and presence of different concentrations of L-Cys. Droplets were prepared with 1.66 μM papain with 10% PEG 8000. Sale bars correspond to 5 μm. (b) Corresponding size distribution histograms with mean droplet sizes.

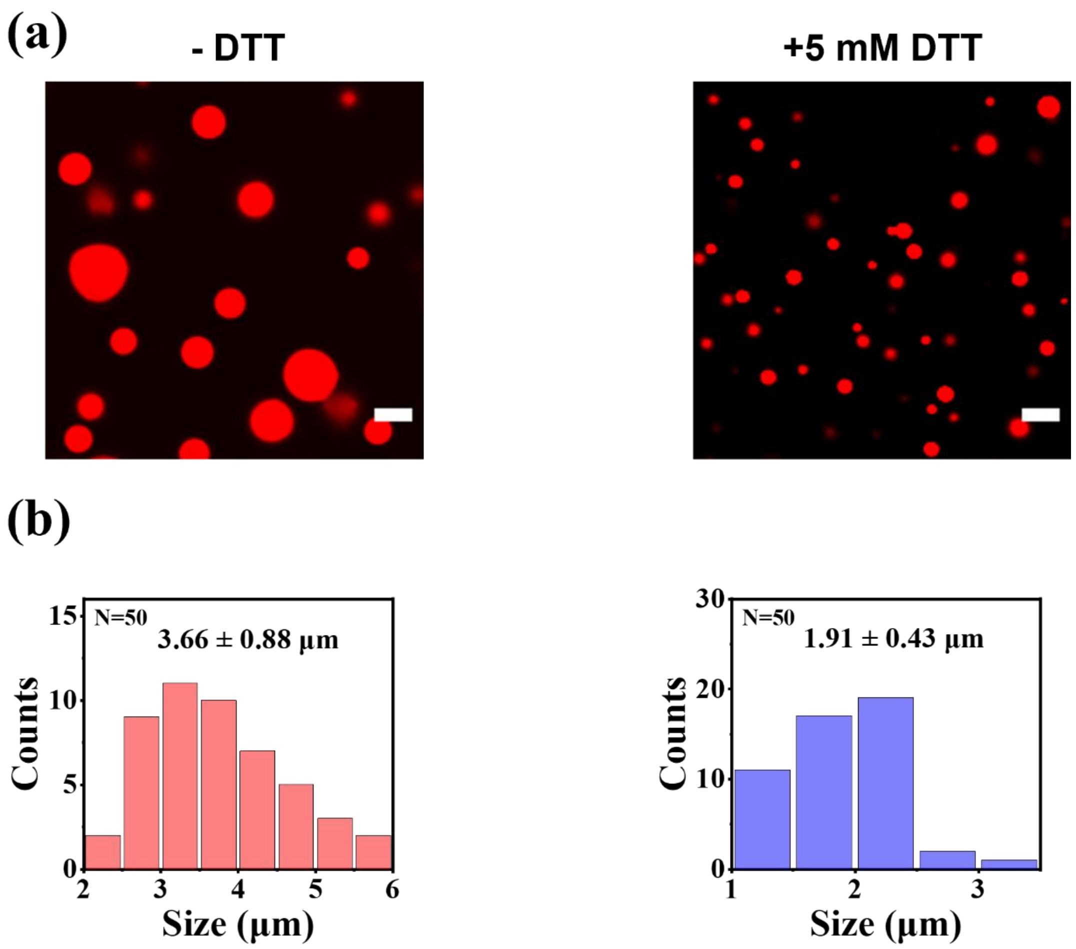


**Figure S21.** (a) Confocal images of RBITC-labeled papain droplets in thew absence and presence of 5 mM dithiothreitol. Droplets were prepared with 1.66 µM papain with 10% PEG 8000. (b) Corresponding size distribution histograms with mean droplet sizes.

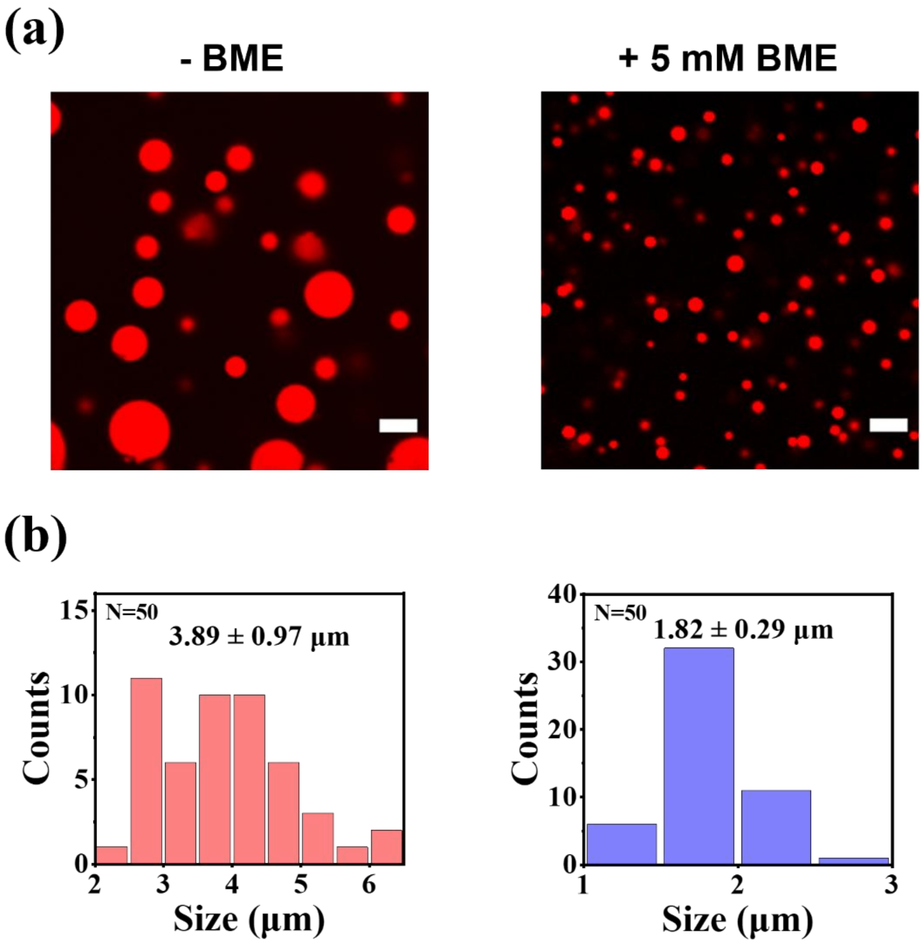


**Figure S22.** (a) Confocal images of RBITC-labeled papain droplets in thew absence and presence of 5 mM $\beta$-mercaptoethanol (BME). Droplets were prepared with 1.66 µM papain with 10% PEG 8000. (b) Corresponding size distribution histograms with mean droplet sizes.

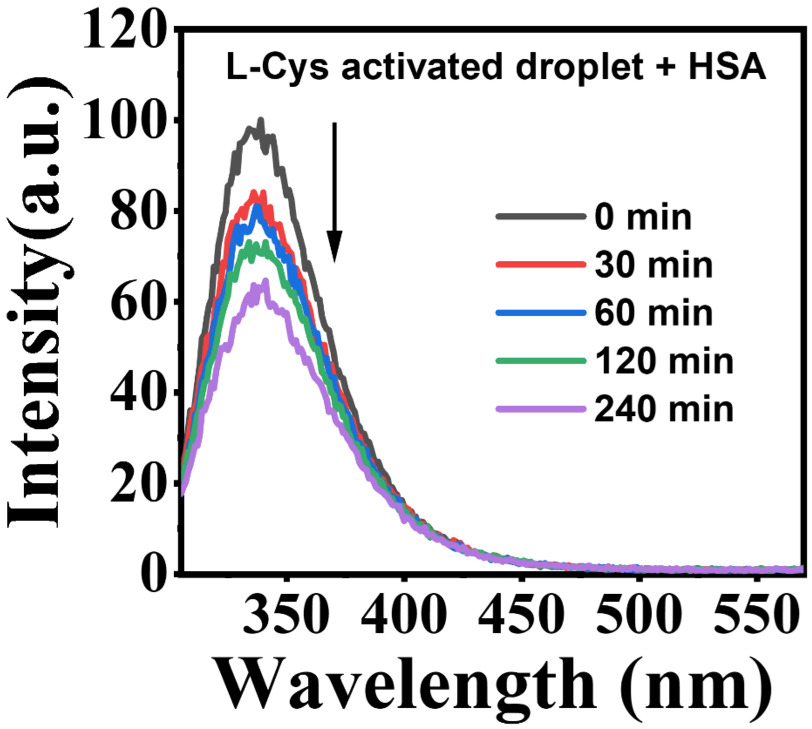


**Figure S23.** Corrected fluorescence spectra ($\lambda_{ex}$ = 295 nm) of 20 μM HSA in the presence of L-Cys activated papain droplets upon 240 min of digestion. Papain droplets were prepared with initial papain concentration of 8 μM in the presence of 10% PEG 8000. These droplets were activated by 5 mM L-Cys.

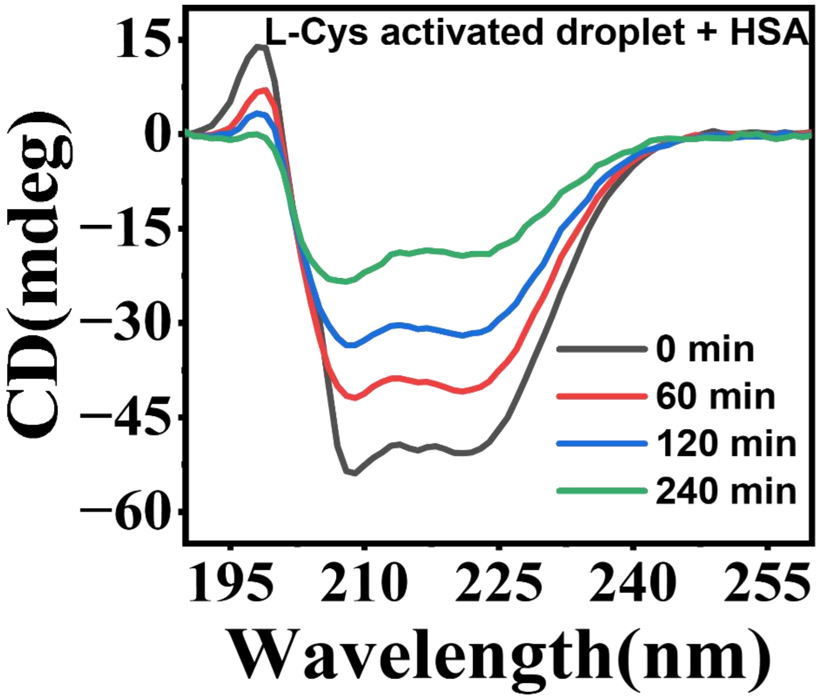


**Figure S24.** Far-UV CD spectra of 4 μM HSA in the presence of L-Cys activated papain droplets over a period of 240 min. Papain droplets were prepared with 1.66 μM papain in the presence of 10% PEG 8000.

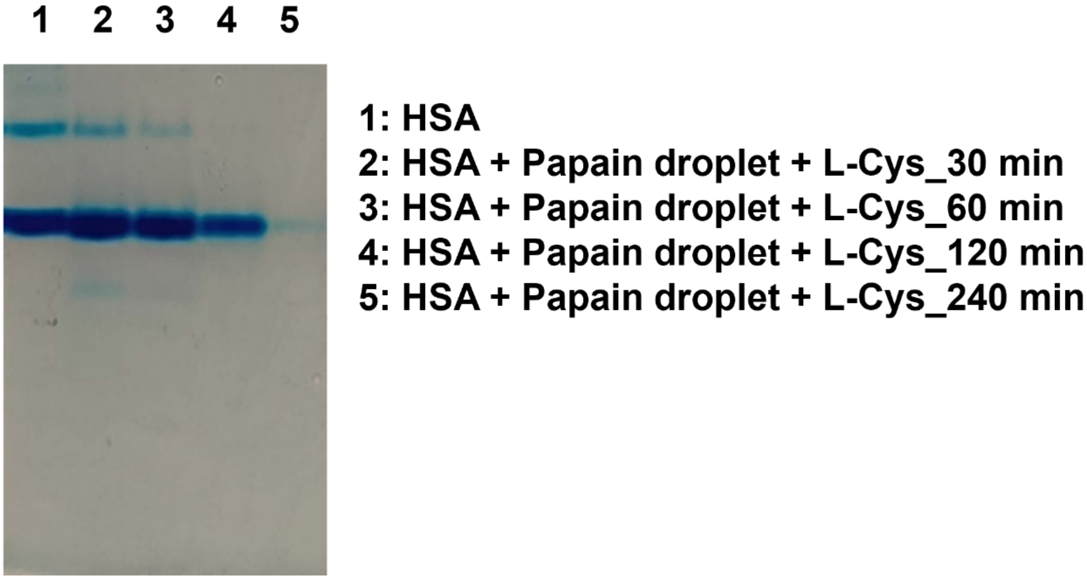


**Figure S25.** Native PAGE data showing the digestion of 20 μM HSA by L-Cys-activated papain droplets over a period of 240 min.

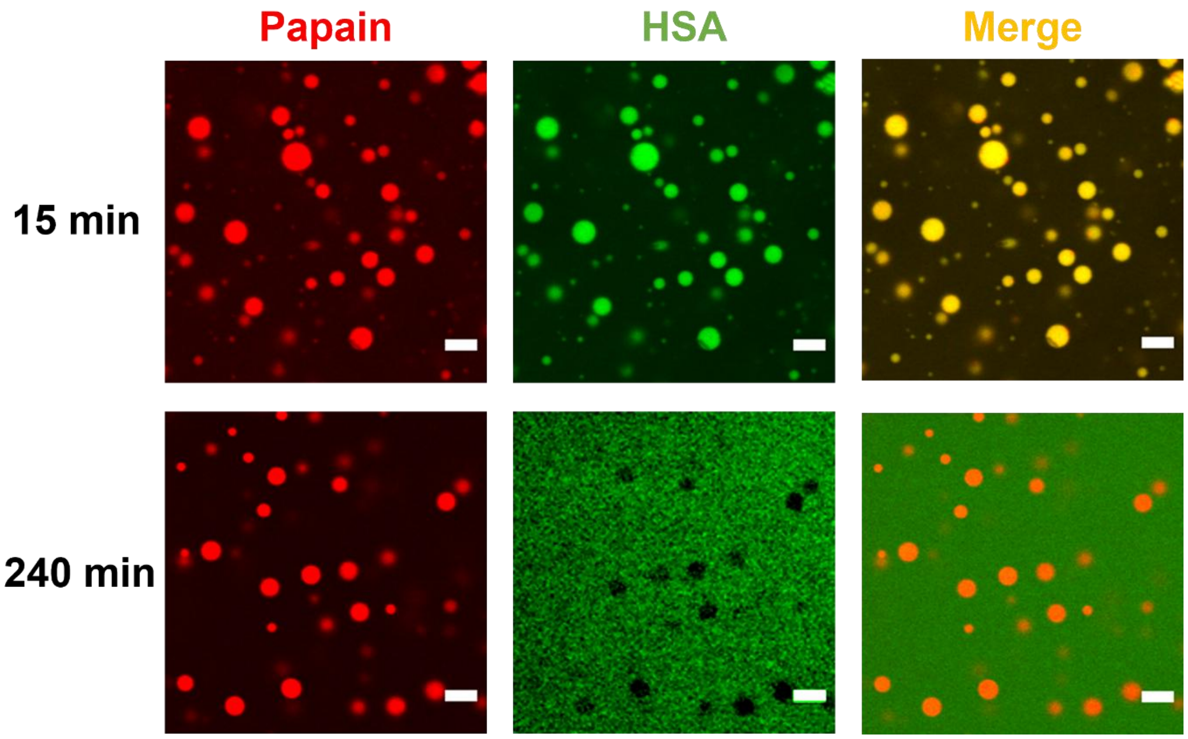


**Figure S26.** Confocal images showing time dependent digestion of FITC-labeled HSA by RBITC-labeled L-Cys-activated papain droplets over a period of 240 min.
